# The long non-coding RNA LINC00941 modulates MTA2/NuRD occupancy to suppress premature human epidermal differentiation

**DOI:** 10.1101/2023.07.25.549662

**Authors:** Eva Morgenstern, Uwe Schwartz, Johannes Graf, Astrid Bruckmann, Markus Kretz

## Abstract

Numerous long non-coding RNAs (lncRNAs) were shown to have functional impact on cellular processes such as human epidermal homeostasis. However, the mechanism of action for many lncRNAs remains unclear to date. Here, we report that lncRNA LINC00941 regulates keratinocyte differentiation on an epigenetic level through association with the NuRD complex, one of the major chromatin remodelers in cells. We find that LINC00941 interacts with NuRD-associated MTA2 in human primary keratinocytes. LINC00941 perturbation changes MTA2/NuRD occupancy at bivalent chromatin domains in close proximity to transcriptional regulator genes, including the *EGR3* gene coding for a transcription factor regulating epidermal differentiation. Notably, LINC00941 depletion resulted in reduced NuRD occupancy at the *EGR3* gene locus, increased EGR3 expression in human primary keratinocytes, as well as increased abundance of EGR3-regulated epidermal differentiation genes in cells and human organotypic epidermal tissue. Our results therefore indicate a role for LINC00941/NuRD in repressing EGR3 expression in non-differentiated keratinocytes, consequentially preventing premature differentiation of human epidermal tissue.

## Introduction

One of the important findings of high-throughput sequencing was that the transcriptional landscape is more complex than originally imagined: It could be shown that only a diminutive fraction of the human genome (< 2%) encode proteins, whereas more than 75% is actively transcribed into non-coding RNAs (ncRNAs), including long non-coding RNAs (lncRNAs) (Djebali *et al,* 2012). LncRNAs are defined as a class of RNAs longer than 200 nucleotides with no or hardly any protein-coding potential (Bonasio & Shiekhattar, 2014). Most lncRNAs are – similar to mRNAs – transcribed by RNA polymerase II, often capped by 7-methyl guanosine (m^7^G) at their 5’ ends, 3’ polyadenylated and spliced (Quinn & Chang, 2016). Many lncRNAs have been shown to be crucial for numerous biological processes, such as X-chromosome inactivation (Strehle & Guttman, 2020), epigenetic control of chromatin (Kazimierczyk & Wrzesinski, 2021) and imprinting (Sanli *et al,* 2018).

The most fundamental function of the human skin and the epidermis, its outermost layer, is to provide a barrier to the external environment. Human epidermis is a stratified surface epithelium constantly renewing approximately every four weeks. This regenerative capacity is mostly maintained by progenitor keratinocytes located in the epidermal basal layer. A subset of daughter cells generated by these progenitor cells dissociate from the basement membrane and migrate to the apical part of the tissue while undergoing a terminal differentiation program. To ensure formation of a functional epidermal barrier, a precise balance between progenitor and differentiated keratinocytes is crucial (Blanpain & Fuchs, 2006; Blanpain & Fuchs, 2009). Several lncRNAs were previously shown to have functional roles in normal epidermal homeostasis including ANCR, TINCR and SMRT-2 (Kretz *et al,* 2012; Kretz *et al,* 2013; Lee *et al,* 2018). Another lncRNA involved in this process is LINC00941, which was demonstrated to be enriched in undifferentiated progenitor keratinocytes and functioned as repressor of keratinocyte differentiation (Ziegler *et al,* 2019). The mechanism by which LINC00941 prevents the onset of epidermal differentiation has not been investigated yet.

Here we show that lncRNA LINC00941 binds several components of the Nucleosome Remodeling and Deacetylase (NuRD) complex, including MTA2, one of its core subunits. Through interaction with NuRD-associated MTA2, LINC00941 modulates occupancy of the epigenetic regulator to mainly transcriptionally repressed and bivalent chromatin sites, thus helping to regulate its gene-regulatory function. Chromatin immunoprecipitation (ChIP) and RNA sequencing approaches showed LINC00941/MTA2/NuRD-mediated repression of EGR3, a transcription factor responsible for keratinocyte differentiation (Kim *et al,* 2019). Thus, LINC00941 prevents premature differentiation by interacting with the MTA2/NuRD complex inhibiting transcription of early and late differentiation genes through their upstream regulator EGR3.

## Results

### The long non-coding RNA LINC00941 interacts with components of the NuRD complex

LINC00941, a lncRNA with partial nuclear localization, was recently shown to regulate expression of early and late key differentiation genes in human epidermis. These included a majority of genes within the small proline-rich (SPRR) protein- and late cornified envelope (LCE) protein-clusters located in the epidermal differentiation complex (EDC) (Ziegler *et al,* 2019). Based on these findings, we hypothesized a LINC00941-mediated regulation of gene expression on a global level through an epigenetic mechanism. To test this hypothesis and to get a deeper insight into the mode of action of LINC00941 in human epidermal homeostasis, we performed RNA-protein interactome analyses to identify proteins associated with LINC00941. For this, RNA pull-down and mass spectrometry (MS) analysis with *in vitro* transcribed, biotinylated LINC00941, as well as cell lysates from primary keratinocytes were performed. Correspondingly, we found that LINC00941 interacts with proteins attributed to chromatin-associated functions, such as nucleosomal DNA binding or histone deacetylase binding properties, as shown by chromatin-associated Gene Ontology (GO) term analysis (Figure 1A). Interestingly, LINC00941-bound proteins were enriched for components of chromatin remodeling complexes – in particular the NuRD complex. A more detailed analysis of the NuRD components identified with our analysis, showed that LINC00941 interacted with CHD4, MTA2, HDAC2, GATAD2A, GATA2DB, RBBP4 and RBBP7, respectively (Figure 1B; Supplement Figure 1; Supplemental Material 1). The NuRD complex combines both, ATP-dependent chromatin remodeling and histone deacetylase activities modulating gene expression. Interestingly, some components of the NuRD complex such as HDAC1/2, CHD4 and MTA2 have previously been demonstrated to fulfill essential functions in murine epidermal development. (Leboeuf *et al,* 2010; Kashiwagi *et al,* 2007; Lu *et al,* 2008). Therefore, we hypothesized that NuRD-mediated epigenetic regulation of differentiation gene expression – including genes in the EDC – might be at least in part controlled by LINC00941. To test our hypothesis, interaction between LINC00941 and the NuRD complex was subsequently further verified. Since multiple components of the NuRD complex such as RBBP4, RBBP7 and HDAC2, can also function as subunits of other multi-protein complexes, including Sin3, Extra Sex Combs/Enhancer of Zeste (ESC/E(Z)), and Nucleosome Remodelling Factor (NuRF) complex (Pantier *et al,* 2017; Dubey *et al,* 2017; Zahid *et al,* 2021), we focused our experiments on MTA2, a core subunit of the NuRD complex (Low *et al,* 2020). The binding of *in vitro* transcribed, biotinylated LINC00941 and NuRD-associated MTA2 from cell lysate with subsequent RNA pull-down was confirmed by Western Blot analysis (Figure 1C). Correspondingly, RNA-immunoprecipitation (RNA-IP) and subsequent qRT-PCR analysis also verified interaction between overexpressed LINC00941 and endogenous MTA2/NuRD in primary keratinocytes (Figure 1D), strongly indicating association of LINC00941 with the chromatin remodeling complex NuRD. This finding suggested a role for the lncRNA to regulate epidermal homeostasis through modulation of NuRD activity or recruitment.

**Figure 1:**
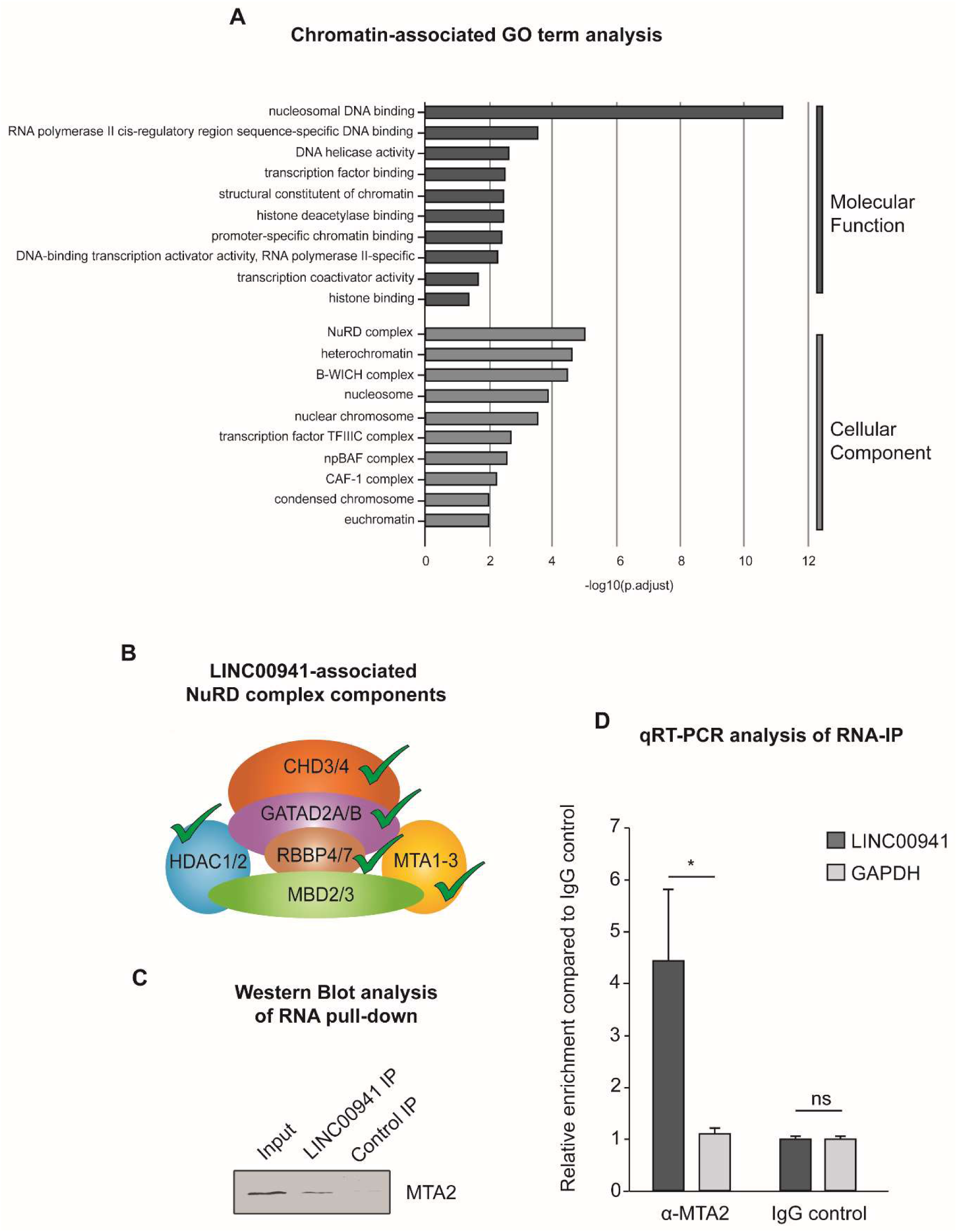
LINC00941 interacts with components of the NuRD complex. A) Chromatin-associated Gene Ontology (GO) terms of LINC00941-bound proteins obtained by mass spectrometry (MS) analysis. The NuRD complex showed the strongest signal with regard to cellular component. B) Scheme of NuRD components found to interact with lncRNA LINC00941 in MS analysis. C) RNA-pull down with *in vitro* transcribed and biotinylated LINC00941 showed interaction with MTA2 in Western Blot analysis. D) RNA-immunoprecipitation (RNA-IP) with overexpressed LINC00941 verified interaction between LINC00941 and MTA2 by qRT-PCR (*n* = 3). Data are presented as ± standard deviation. Statistical significance was tested by an unpaired *t*-test and corrected for multiple testing after Bonferroni (*adj. *P*-value < 0.05 and ns = not significant).

### NuRD-associated MTA2 prevents premature differentiation of human keratinocytes

We recently reported that LINC00941 was highly induced in undifferentiated progenitor keratinocytes and reduced in abundance as keratinocyte differentiation progresses (Ziegler *et al,* 2019). To test if NuRD-associated MTA2 shows a similar dynamic regulation during human keratinocyte differentiation, we performed qRT-PCR analysis of MTA2 throughout six time points of calcium-induced, human keratinocyte differentiation. We found that *MTA2* mRNA was highly abundant in non- and poorly differentiated keratinocytes but repressed during keratinocyte differentiation, thus exhibiting an expression pattern similar to LINC00941 (Figure 2A). Correspondingly, Western Blot analyses with MTA2-specific antibodies also confirmed declining protein amounts of MTA2 in calcium-induced, differentiated keratinocytes (Figure 2B). These results of correlating expression patterns further suggested a functional interaction between the LINC00941 and the NuRD complex.

**Figure 2:**
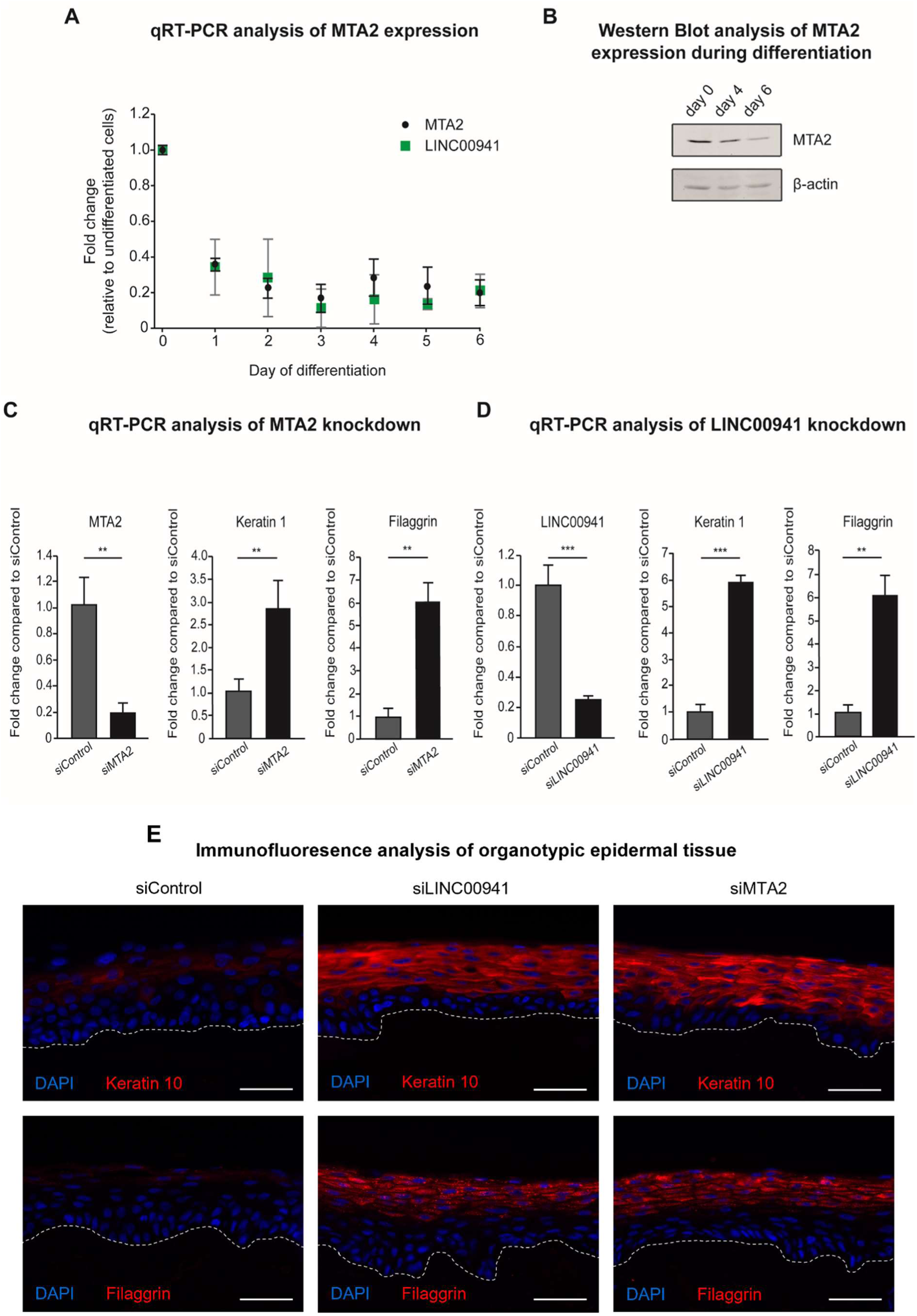
NuRD-associated MTA2 prevents premature onset of keratinocyte differentiation. A) qRT-PCR analysis of LINC00941 and MTA2 repression during calcium-induced differentiation in primary keratinocytes compared to undifferentiated keratinocytes (day 0) (*n* = 3-4). Data are presented as mean ± standard deviation. B) Western Blot analysis of MTA2 repression during calcium-induced differentiation in primary keratinocytes (day 4 and day 6) compared to undifferentiated keratinocytes (day 0). C,D) qRT-PCR analysis of either MTA2 or LINC00941 knockdown. SiPool-mediated knockdown of MTA2 (C) and LINC00941 (D), respectively, resulted in increased abundance of early and late differentiation marker keratin 1 and filaggrin on day 3 of differentiation in organotypic epidermal tissue (*n* = 3-5 tissue cultures/knockdown group). Data are presented as mean ± standard deviation. Statistical significance was tested by an unpaired *t*-test and corrected for multiple testing after Bonferroni (**adj. *P*-value < 0.01, ***adj *P*-value < 0.001). E) Immunofluorescence (IF) analysis showed increased level of early and late differentiation proteins keratin 10 and filaggrin. Dashed line indicates the basement membrane, nuclei are shown in blue, and the differentiation proteins keratin 10 and filaggrin are shown in red (*n* = 3-5 tissue cultures/knockdown group, one exemplary picture for each group is depicted). Scale bar: 100 µm.

To test whether MTA2 affects human epidermal homeostasis in a similar manner as LINC00941, we generated MTA2-deficient organotypic human epidermis using pools of 11-30 siRNAs to achieve efficient and specific MTA2 depletion. MTA2-deficient organotypic epidermis showed increased mRNA abundance of the early differentiation gene keratin 1 and the late differentiation gene filaggrin (Figure 2C). Correspondingly, LINC00941-deficient organotypic epidermis yielded a similarly increased abundance of both mRNA populations (Figure 2D). Therefore, both LINC00941- or MTA2-deficient tissues showed premature and more advanced differentiation compared to control-treated organotypic epidermis, as indicated by increased protein abundances of keratin 10, an early differentiation marker co-expressed with keratin 1, and the late differentiation marker filaggrin (Figure 2E). The above results suggested a common role for the complex consisting of LINC00941 and NuRD in repressing a premature onset of keratinocyte differentiation in human epidermal tissue.

### MTA2/NuRD occupies regulatory regions in keratinocytes

Since LINC00941 interacted with the NuRD complex and both appeared to have overlapping functionality, we hypothesized a role for LINC00941 modulating NuRD-mediated epigenetic regulation of genes relevant for epidermal homeostasis. To examine the genome-wide impact of LINC00941 deficiency on NuRD binding to chromatin, we performed ChIP sequencing with LINC00941-deficient and control-treated keratinocytes using antibodies directed against MTA2. First, we wanted to get a deeper understanding of the general chromatin binding behavior of the NuRD-complex in keratinocytes and therefore identified genome-wide MTA2/NuRD binding sites (*n* = 3,613) (Supplemental Material 2), which were highly reproducible between replicates (Supplement Figure 3A). We observed enrichment of MTA2/NuRD occupancy at promoters (Figure 3A; Supplement Figure 3B). GO term analysis of NuRD-associated promoters included genes regulating cell fate commitment as well as morphogenic and developmental processes (Figure 3B; Supplement Figure 3C). Importantly, these terms also comprise targets important for keratinocyte development and differentiation such as MAFB, EPHA2 and SOX9.

**Figure 3:**
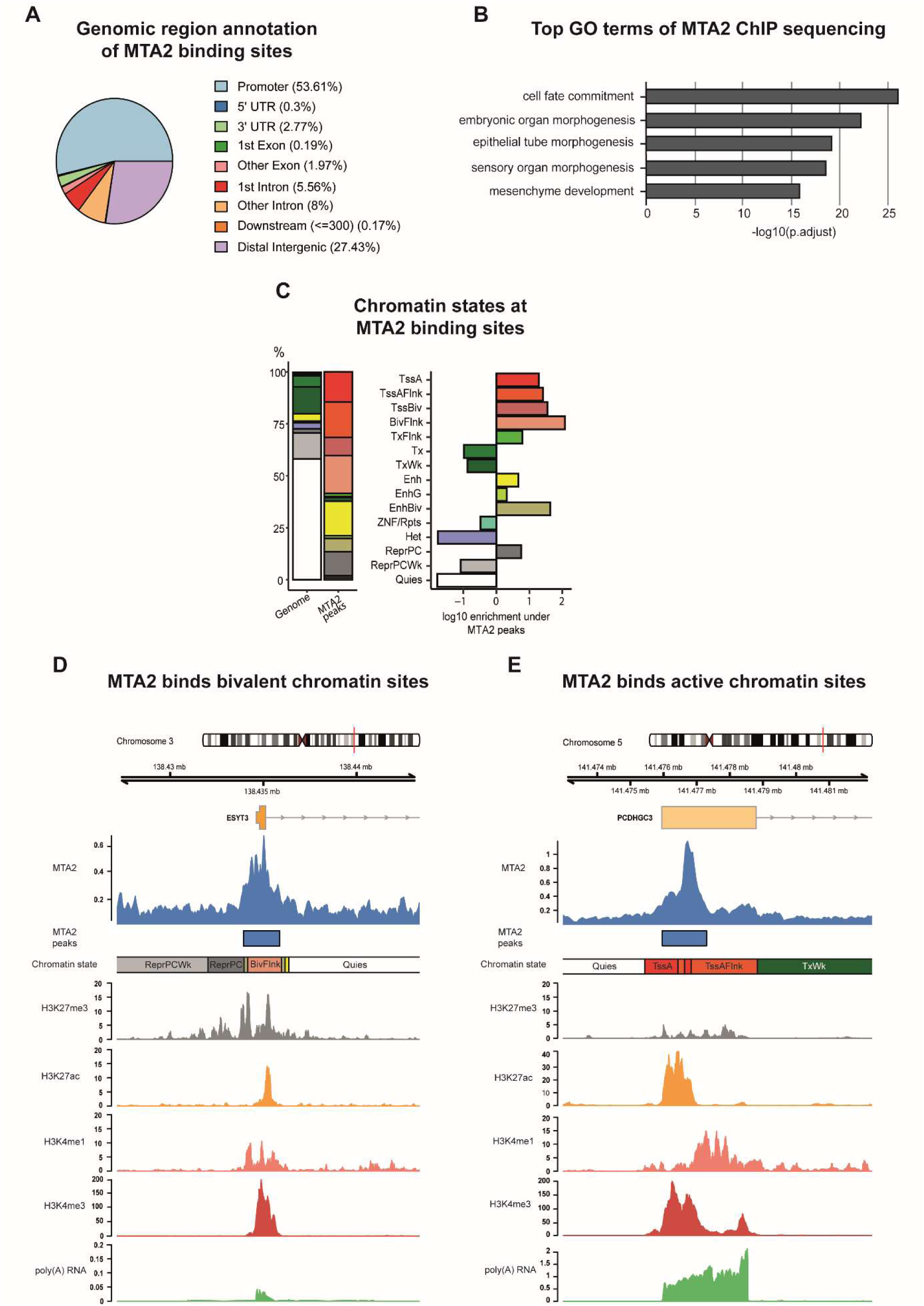
MTA2/NuRD occupies regulatory regions in keratinocytes. A) Pie chart showing genomic region annotation of MTA2 binding sites, preferentially binding to promoter and distal intergenic regions. The genome annotation of protein-coding genes was used. The promoter region was defined as +/- 1000 bp distance to the TSS. B) Top GO terms at MTA2 binding sites. The respective GO terms also included genes involved in keratinocyte differentiation. GO term analysis was restricted to terms associated with a minimum of 250 and a maximum of 350 genes. C) Bar plots showing the distribution of primary keratinocyte chromatin states at MTA2 binding sites. The left bar represents the distribution of chromatin states over the whole genome and the right bar at the MTA2 binding sites. The enrichment of the chromatin states at MTA2 binding sites is shown as the log10 fold-change of the respective chromatin state against the whole genome distribution. MTA2 was found to bind activated (TssA, TssAFlnk, Enh, EnhG), repressed (ReprPC) and bivalent (TssBiv, EnhBiv, BivFlnk) chromatin states. Abbreviations: Tss = Transcription Start Site; A = Active; Flnk = Flanking; Biv = Bivalent; Tx = Transcription; Wk = Weak; Enh = Enhancer; G = Genic; Rpts = Repeats; Het = Heterochromatin; Repr = Repressed; PC = Polycomb; Quies = Quiescent D, E) Genome browser view of MTA2 binding sites at selected genomic regions. Tracks of chromatin states, histone modifications, and transcription in primary keratinocytes obtained from roadmap (accession: E057) are shown below the MTA2 ChIP sequencing tracks. Abbreviations: Tss = Transkription Start Site; A = Active; Flnk = Flanking; Biv = Bivalent; Tx = Transkription; Wk = Weak; Quies = Quiescent

To analyze the chromatin context at MTA2/NuRD occupied sites, we used the epigenome roadmap chromatin state model of primary keratinocytes (Kundaje *et al,* 2015) and found that NuRD-associated MTA2 bound to various chromatin states at regulatory elements (Figure 3C). MTA2/NuRD was not only associated with activated chromatin states such as active transcription start sites (TssA, TssAFlnk) and enhancers (Enh, EnhG) but also with repressed chromatin states (ReprPC). Furthermore, it was also frequently bound to bivalent chromatin states including bivalent transcription start sites, enhancers, and related flanking regions (TssBiv, BivFlnk and EnhBiv). A more detailed analysis of some representatives of those different chromatin states confirmed the versatile chromatin binding behavior of MTA2/NuRD: ESYT3, a membrane-associated protein predicted to bind calcium ions and phospholipids, represented one example for a bivalent MTA2/NuRD occupancy site. It harbored both, active (H3K4me3, H3K27ac and H3K4me1) and repressive (H3K27me3) histone marks at the MTA2 occupied site, resulting in sparse expression of the target gene. The *PCDHGC3* gene locus represents a typical example for MTA2/NuRD occupancy within activated chromatin sites (TssA, TssAFlnk). Correspondingly, solely activating histone marks could be detected (Figure 3E). PCDHGC3 is an adhesion protein, potentially calcium-dependent and expressed in human keratinocytes. The broad range of MTA2/NuRD-associated chromatin states in human keratinocytes emphasized a complex role of NuRD as an epigenetic regulator complex – beyond the role as transcriptional repressor (Miccio *et al,* 2010; Hu & Wade, 2012; Pundhir *et al,* 2023). Interestingly, its distribution of differential peaks at genomic features of primary keratinocytes showed a stronger preference for promoter regions opposite to several reports published previously, which detected a preference for intragenic regions and a subordinate role of promoters (Lu *et al,* 2019; Arends *et al,* 2019; Marques *et al,* 2020). However, ChIP sequencing analysis of NuRD-associated MBD3 in mouse embryonic stem cells found a similar enrichment of the NuRD complex at promoter regions, generally indicating a cell-type and -state dependent occupancy pattern (Yildirim *et al,* 2011).

### LINC00941 dependency of MTA2/NuRD binding at *EGR3*

Next, we tested whether the binding behavior of MTA2/NuRD changes in LINC00941-deficient cells. Unsupervised principal component analysis (PCA) revealed that the highest variance in MTA2/NuRD occupancy across all samples could be explained by the siRNA-mediated LINC00941 knockdown, thus suggesting a regulatory role of LINC00941 on MTA2/NuRD chromatin binding (Figure 4A). Differential occupancy analysis resulted in 33 significantly altered MTA2/NuRD binding sites upon LINC00941 depletion (FDR threshold of 5%) (Figure 4B). 67% of all differential MTA2/NuRD occupied sites were located directly at the TSS, suggesting LINC00941 as putative regulator of transcription through MTA2/NuRD (Supplement Figure 4A). Most of the differential MTA2/NuRD occupied sites were marked by the repressive histone modification H3K27me3 corresponding to Polycomb repressed and bivalent chromatin states (Supplement Figure 4B), which is in agreement with the known function of NuRD to enhance PRC2 binding (Kim *et al,* 2015). In line with that, most of the associated genes were not or sparsely transcribed in undifferentiated keratinocytes (Figure 4B). The majority of 29 target sites showed significantly reduced occupancy of MTA2/NuRD in LINC00941 knockdown cells. In these cases, LINC00941 appeared to facilitate NuRD complex binding to the target genes, thereby promoting transcriptional repression by PRC2. Interestingly, four binding sites were associated with active chromatin states and their corresponding target genes (*AP3D1*, *MROH6*, *SLC25A45* and *C2CD2*) were considerably expressed in undifferentiated keratinocytes. Here, LINC00941 deficiency resulted in enhanced MTA2/NuRD binding indicating that LINC00941 bound to the MTA2/NuRD complex appeared to diminish binding of the NuRD complex to these target genes and ensures their transcription. Furthermore, many of the genes associated with altered MTA2/NuRD occupancy in LINC00941-deficient keratinocytes were transcriptional regulators, some of which were known to be involved in differentiation processes. Therefore, we hypothesized that LINC00941/NuRD might suppress the expression of transcriptional activators. This repression would be restricted to non- and weakly differentiated strata of the epidermis, since the levels of LINC00941 and MTA2, respectively, declined during differentiation (Figure 2A). To verify this hypothesis, we analyzed to what extend MTA2/NuRD-associated genes dynamically changed their expression during keratinocyte differentiation and whether their transcription is influenced by LINC00941 knockdown (Figure 4C). In addition to a solute carrier protein (SLC25A45) (Supplement Figure 4C), only the transcription factor EGR3 was fulfilling both criteria: *EGR3* showed both significant upregulation upon LINC00941 knockdown in organotypic epidermal tissue and upregulation during keratinocyte differentiation. Interestingly, an analysis of the chromatin conformation states at the *EGR3* gene locus revealed that it was annotated as a bivalent and repressed site up to and including day 2.5 of calcium-induced differentiated keratinocytes (Figure 4D). No later than day 5.5, this site was associated with an active chromatin state. These findings were also supported by ChIP sequencing data of histone modifications and RNA sequencing data of calcium-induced differentiated keratinocytes from ENCODE. Up to day 2.5 of differentiation, repressive H3K27me3 was the most prevalent mark in undifferentiated keratinocytes but was erased in course of differentiation. On the other hand, the transcription-activating chromatin modification H3K27ac increased over time. Correspondingly, the first *EGR3* transcripts were detectable at day 5.5 based on RNA sequencing data in wild-type keratinocytes (Figure 4C and 4D). This led us hypothesize that LINC00941/ NuRD binds to the *EGR3* gene locus during repressed and bivalent states, respectively, promoting transcriptional repression. In the course of the differentiation process, the chromatin states at the *EGR3* locus changed and LINC00941 as well as MTA2 were transcriptionally repressed themselves (Figure 2A), thus allowing for active transcription of *EGR3*. In accordance with this hypothesis, LINC00941 knockdown led to premature increase of EGR3 abundance in organotypic epidermis, thus accelerating this tightly regulated process (Figure 4E). In conclusion, our data strongly suggested a LINC00941-regulated, epigenetic repression of the *EGR3* gene locus by MTA2/NuRD, therefore preventing premature transcriptional activity during epidermal homeostasis.

**Figure 4:**
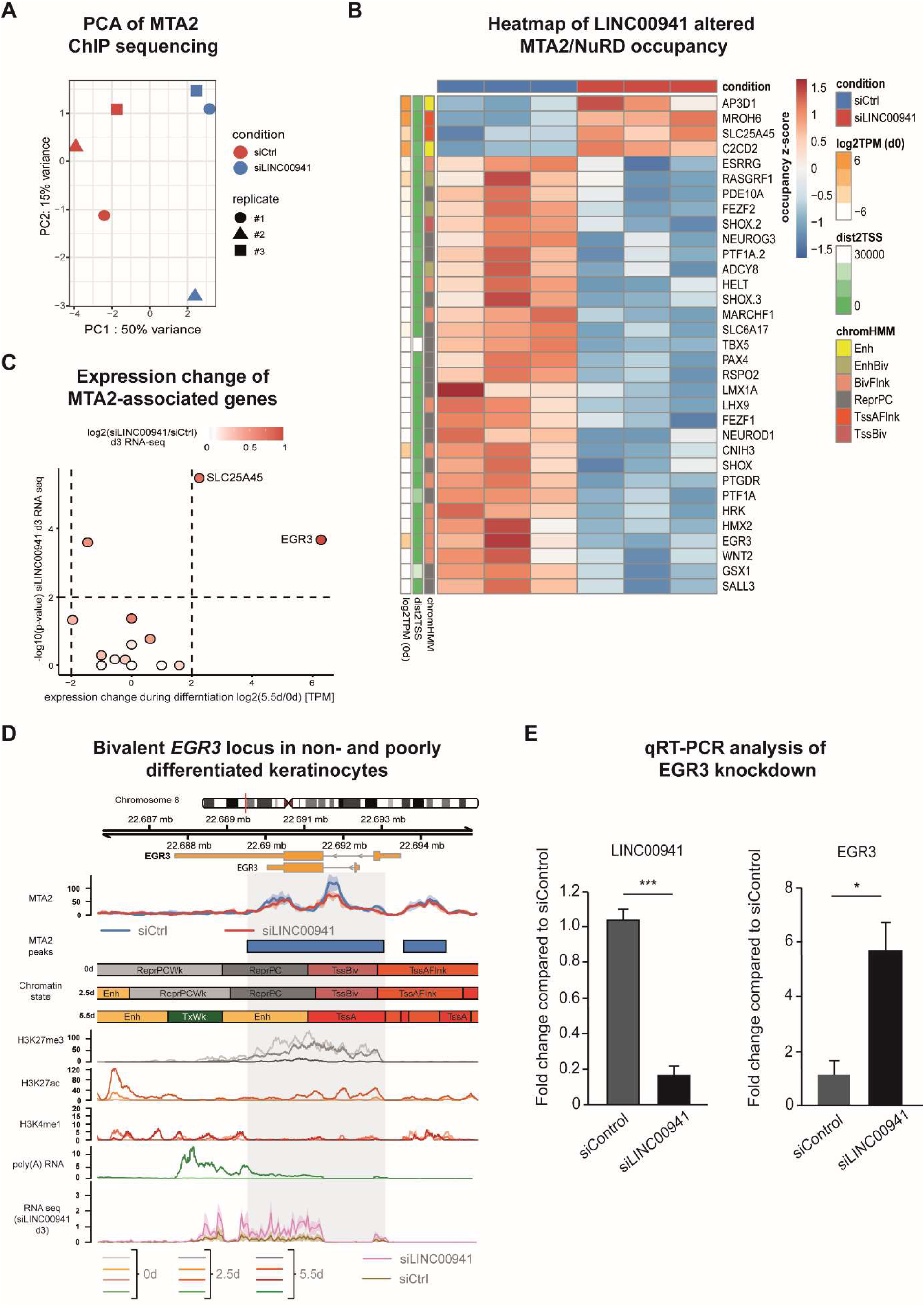
LINC00941 dependency of NuRD-associated MTA2 binding in *EGR3*. A) Principal Component Analysis (PCA) of MTA2 ChIP sequencing experiments upon LINC00941 knockdown. B) Heatmap illustrating differential MTA2 bound sites (*n* = 33 at FDR of 5%) upon LINC00941 knockdown. MTA2/NuRD showed different chromatin occupancy in LINC00941-deficient cells compared to control. Z-scores of normalized fragment counts under differential MTA2 binding sites are indicated by the color code. Row labels on the right depict the gene nearest to the differential MTA2 binding site. Associated chromatin state (chromHMM) and distance to (dist2TSS) or expression level (log2(TPM) at 0d) in primary keratinocytes of associated genes are indicated on the left side. Abbreviations of chromatin states: Tss = Transcription Start Site; A = Active; Flnk = Flanking; Biv = Bivalent; Enh = Enhancer; Repr = Repressed; PC = Polycomb. C) Scatter plot illustrating expression change in keratinocyte differentiation or upon LIN00941 knockdown of MTA2-associated genes. EGR3 showed the strongest changes upon both keratinocyte differentiation and LINC00941 knockdown. The x-axis represents the log2 fold change in expression after 5.5d of calcium-induced keratinocyte differentiation against primary keratinocytes. TPM values were obtained from ENCODE Portal (0d: ENCFF423MWU; 5.5d: ENCFF379PNP). The y-axis denotes the -log10 transformed p-values derived from differential gene expression analysis between cells treated with siLINC00941 and control keratinocytes after 3 days of differentiation induction (Ziegler et al., 2019). The log2 fold change in expression upon LINC00941 knockdown is indicated by shades of red. Genes exhibiting a p-value < 0.01 and |log2 fold change| > 2 were highlighted. D) Genome browser view of differential MTA2 binding sites at *EGR3* locus. EGR3 showed bivalent chromatin states in non- and poorly differentiated keratinocytes. Tracks of chromatin states, histone modifications, and transcription in primary undifferentiated (0d) and calcium-treated differentiated (2.5d, 5.5d) keratinocytes obtained from ENCODE portal are shown below the MTA2 ChIP sequencing tracks. Color shades in H3K27me3, H3K27ac, H3K4me1 and RNA sequencing tracks indicate the time points of differentiation: light shade for 0d, medium shade for 2.5d and dark shade for 5.5d. The bottom track shows the RNA sequencing coverage of siLINC00941 transfected (pink) and control (brown) keratinocytes after 3d of differentiation. The dashed box highlights the differential MTA2 binding site at EGR3. Abbreviations: Tss = Transcription Start Site; A = Active; Flnk = Flanking; Biv = Bivalent; Tx = Transcription; Wk = Weak; Enh = Enhancer; Repr = Repressed; PC = Polycomb; Quies = Quiescent E) siPool-mediated knockdown of LINC00941 resulted in decreased *EGR3* level in organotypic epidermal tissue (*n* = 3-5 tissue cultures/knockdown group). Data are presented as mean ± standard deviation. Statistical significance was tested by an unpaired t-test and corrected for multiple testing after Bonferroni (*adj. P-value < 0.05, ***adj. P-value < 0.001).

### EGR3 promotes keratinocyte differentiation

EGR3 was recently described as regulator of keratinocyte differentiation (Kim *et al,* 2019). In addition to differentiation genes, Kim et al. identified 20 functional EGR3-regulated genes, which showed increased expression in calcium-induced differentiated keratinocytes, reduced abundance upon EGR3 knockdown and which were found to be co-expressed with *EGR3*. To further support our hypothesis, that LINC00941-dependent decrease of NuRD binding at *EGR3* affected its function, we performed an overlap between RNA sequencing data from LINC00941-deficient cells and EGR3-regulated genes found by Kim et al. Interestingly, all EGR3-regulated genes were increased in abundance in LINC00941 depleted, organotypic epidermal tissues even though they are normally not expressed until day 5.5 in calcium-induced differentiated keratinocytes (Figure 5A; Supplement Figure 5A). To verify the functional impact of EGR3 in our organotypic model for human epidermal homeostasis, we generated organotypic epidermal tissue with EGR3-deficient and control-treated keratinocytes (Figure 5B and 5C). We were able to detect significantly reduced mRNA levels of keratin 1 and filaggrin, respectively. In addition, early and late differentiation markers keratin 10 and filaggrin showed severely reduced abundances in EGR3-deficient organotypic epidermis. We also detected severely reduced abundance of LINC00941-regulated differentiation genes located within the EDC cluster as a consequence of EGR3 knockdown (Supplement Figure 5B). These results indicated an important role for EGR3 in promoting keratinocyte differentiation in human epidermal tissue not only through reduced abundance of EDC-associated late but also early differentiation genes.

**Figure 5:**
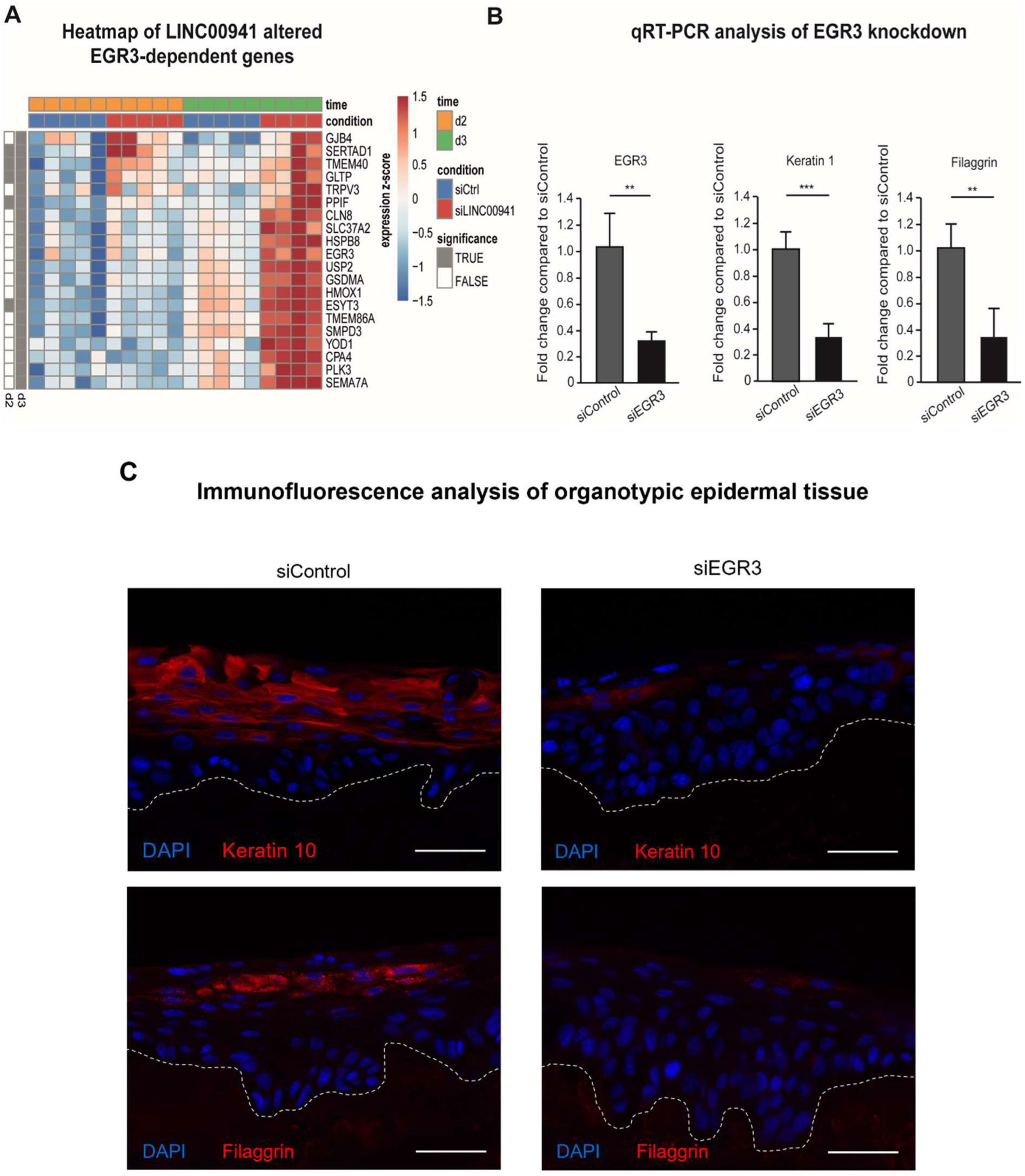
EGR3 induces differentiation of human organotypic epidermis. A) Heatmap showing transcriptional changes of *EGR3* and EGR3-regulated genes in keratinocyte differentiation upon LINC00941 knockdown. Genes regulated by EGR3 in keratinocytes were selected based on the study of Kim et al. (2019). Expression data of siLINC00941 treated keratinocytes after 2d and 3d were obtained from GEO (GSE118077) (Ziegler et al., 2019) and reanalyzed. Significant expression changes (FDR of 5%) of pair-wise comparisons between siLINC00941 and control at each time point are indicated at the left sidebar (grey = significant, white = not significant). B) qRT-PCR analysis of EGR3 knockdown. SiPool-mediated knockdown of EGR3 in primary keratinocytes resulted in decreased abundance of early and late differentiation marker keratin 1 and filaggrin on day 3 of differentiation in organotypic epidermal tissue (*n* = 4-6 tissue cultures/knockdown group). Data are presented as mean ± standard deviation. Statistical significance was tested by an unpaired t-test and corrected for multiple testing after Bonferroni (**adj. *P*-value < 0.01, ***adj. *P*-value < 0.001). C) Immunofluorescence (IF) analysis showed decreased level of early and late differentiation marker keratin 10 and filaggrin. Dashed line indicates the basement membrane, nuclei are shown in red (*n* = 4-6 tissue cultures/knockdown group, one exemplary picture for each group is depicted). Scale bar: 100 µm.

In conclusion, our data strongly suggests a role for lncRNA LINC00941 in regulating recruitment of the chromatin remodeling complex NuRD to target gene loci, thus controlling epidermal homeostasis. Correspondingly, LINC00941/NuRD-dependent epigenetic repression of the transcription factor EGR3 – a regulator of epidermal differentiation – represents a lncRNA-dependent, regulatory mechanism for modulation of the epidermal homeostatic balance.

## Discussion

Previous work identified LINC00941 as negative regulator of premature human keratinocyte differentiation, but the exact mechanism remained unknown (Ziegler *et al,* 2019). LINC00941-mediated regulation of many genes within the SPRR protein-as well as the LCE protein-clusters located within the EDC suggested a role for LINC00941 as an epigenetic regulator of transcription (Ziegler *et al,* 2019; Morgenstern & Kretz, 2023). The above study has confirmed this hypothesis and unraveled the mode of action of LINC00941 as transcriptional modulator of epidermal homeostasis for the first time. This study demonstrated LINC00941-dependent MTA2/NuRD binding at several chromatin sites, including *EGR3,* a transcriptional regulator of keratinocyte differentiation (Figure 4B). We could show epigenetic repression of *EGR3* transcription in undifferentiated progenitor keratinocytes due to MTA2/NuRD-mediated gene silencing, with MTA2/NuRD occupancy being dependent on LINC00941. However, absence of LINC00941-mediated MTA2/NuRD binding to the *EGR3* gene locus facilitated its expression and resulted in increased abundance of its downstream targets involved in keratinocyte differentiation (Figure 4D, 4E, 5 and 6).

Similar to the above study, LINC00941 was previously found to also influence mRNA transcription. However, this preceding study described the involvement of LINC00941 in cis-acting processes: LINC00941 regulates expression of its nearby gene *CAPRIN2* through interacting with CTCF, a transcriptional regulator mediating chromatin looping (Ai *et al*, 2020). Another work revealing a role of LINC00941 as regulator of mRNA transcription was recently published: Lu et al. (2023) described LINC00941-mediated recruiting of ILF2 and YBX1 to *SOX2* promoter region to enhance its transcription. In both cases, LINC00941 was found to have an activating gene regulatory role in both oral squamous and esophageal cell carcinoma, respectively. Here, interaction of LINC00941 with the NuRD complex could be demonstrated. Predominantly acting as transcriptional repressor in non-cancerous epithelial cells and tissue bound to the NuRD complex, LINC00941 is thus another lncRNA that plays an important role in epidermal homeostasis, in addition to the lncRNAs ANCR, TINCR, SMRT-2, BLNCR and PRANCR (Kretz *et al*, 2012; Kretz *et al*, 2013; Lee *et al*, 2018; Tanis *et al*, 2019; Cai *et al*, 2020).

We previously showed that LINC00941 regulated genes associated with the process of epidermal stratification and differentiation – including genes located within the epidermal differentiation cluster (Ziegler *et al*, 2019). Interestingly, our results presented above did not reveal direct occupancy of the NuRD complex at genes within the EDC. Instead, we found LINC00941-controlled MTA2/NuRD occupancy at gene loci of various transcription factors, including *EGR3*. Correspondingly, LINC00941 likely acts through epigenetic repression of EGR3, a transcriptional regulator of differentiation gene expression. Our data indicate that *EGR3* transcription was likely repressed in non- and poorly differentiated strata of human epidermis through binding of LINC00941/NuRD. Since MTA2/NuRD and LINC00941 abundance decreased upon differentiation (Figure 2A), epigenetic repression of *EGR3* transcription was reduced (Figure 4D). This likely resulted in EGR3-mediated transcription of downstream targets, including early and late differentiation genes – many of which are located within the EDC and induced in LINC00941-deficient epidermis – as demonstrated by the analyses presented here (Figure 6) as well as others (Kim *et al*, 2019).

**Figure 6:**
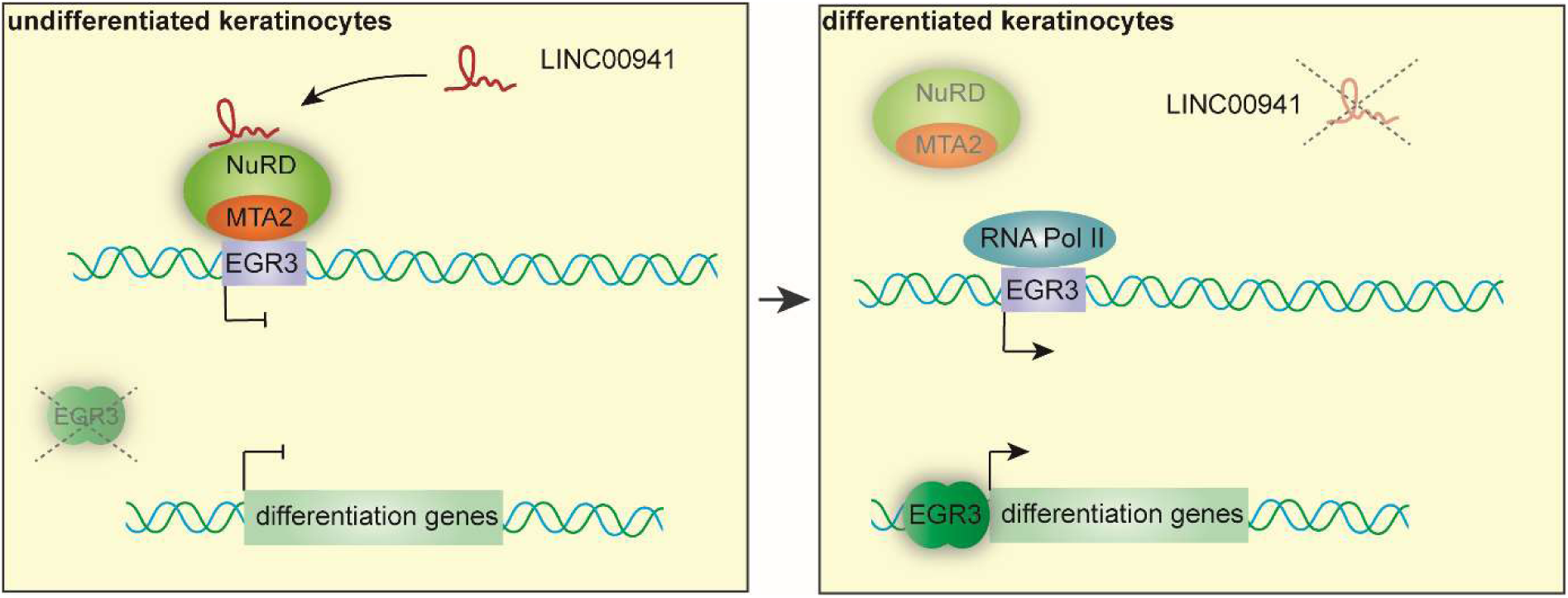
LINC00941 regulates epidermal differentiation genes through modulation of MTA2/NuRD at the *EGR3* gene locus. Predicted mechanism of LINC00941. Interaction of LINC00941 with the MTA2/NuRD complex modulates its binding to chromatin, thus repressing *EGR3* expression in non- and poorly differentiated keratinocytes. During keratinocyte differentiation, LINC00941 and MTA2 abundances decrease and *EGR3* is actively transcribed, which in turn results in expression of its downstream differentiation gene targets and consequentially acceleration of keratinocyte differentiation.

As part of the above study, we found an enrichment of LINC00941/MTA2/NuRD occupancy to bivalent and repressed chromatin states, respectively, including the *EGR3* gene locus (Figure 4B). Bivalent chromatin states were first described by Bernstein et al (2006) as they discovered nucleosomes marked with both repressing H3K27me3 and activating H3K4me3 histone modifications at the same time. In mouse embryonic stem (ES) cells, they found these bivalent domains frequently in close proximity to non- and poorly expressed developmental transcription factor genes and other genes important for development, that were poised for subsequent activation during ES cell differentiation. However, these repressed regions resolved during ES cell differentiation (Bernstein *et al*, 2006). Therefore, it was initially proposed that bivalent histone modifications keep developmental genes in a silenced but translational state allowing either rapid activation or stable silencing during ES cell differentiation. In the later course this definition was revised since bivalent domains were not only discovered in mouse and human ES cells but also in mammalian adult stem cells and adult tissues including keratinocytes (Kinkley *et al*, 2016; Mikkelsen *et al*, 2007; Barrero *et al*, 2013). Consequently, bivalency is now commonly seen as a way of expression fine-tuning during cell development and cell fate decisions, respectively, to prevent unscheduled gene activation providing robustness and plasticity but reduced noise during repression (Voigt *et al*, 2013). One of the central components in establishing and maintaining bivalency are the PcG proteins PRC1 and PRC2 (Harikumar & Meshorer, 2015), which are thought to be recruited by lncRNAs among others (Voigt *et al*, 2013). In case of the NuRD complex, its role in interrelation with PcG proteins is better explored. It could be demonstrated that this chromatin remodeler deacetylates H3K27 in ES cells enabling PcG protein recruitment. This in turn, can catalyze trimethylation of H3K27 at NuRD target genes allowing cell differentiation (Kaji *et al*, 2006; Reynolds *et al*, 2012; Hu & Wade, 2012; Kim *et al*, 2015). Consequently, the NuRD complex controls the sensitive equilibrium between acetylation and methylation of histones, thus influencing the expression of genes critical for cell fate decisions. These findings are in accordance with our results since we also detected MTA2/NuRD at active, repressed, and bivalent chromatin states (Figure 3C). For the first time, however, we could show that the MTA2/NuRD binding at bivalent domains in close proximity to transcriptional regulators, such as the transcription factor gene *EGR3*, depend on the presence of a lncRNA. Therefore, we hypothesize that LINC00941 might directly influence the maintenance of bivalent regions as it recruits NuRD. The resulting repression of EGR3 is normally maintained in non- and poorly differentiated keratinocytes until this bivalent site is resolved through external factors of unknown origin. As LINC00941- and NuRD-associated MTA2 abundance decreased upon progressive differentiation, they were supposed to contribute to resolving bivalent domains. This assumption was further supported by the fact that LINC00941-deficient cells showed premature expression of *EGR3* coding for a transcription factor important for keratinocyte cell fate with regard to differentiation. This implied an early removal of the bivalent status. The precise mechanism of NuRD complex recruitment and modulation of activity, respectively, is not known to date and represents a matter of ongoing research. However, recent studies have revealed CHD4 as actual RNA-binding subunit of NuRD, thus tethering ncRNAs to target chromatin sites mediating DNA-RNA triplexes (Zhao *et al*, 2018; Ullah *et al*, 2022). In summary, our findings revealed the mode of action of LINC00941 involved in epidermal homeostasis. This process was dependent on LINC00941-mediated recruitment of NuRD to bivalent chromatin sites, which could also be found at the *EGR3* gene locus. LINC00941/NuRD regulated repression of EGR3 gene expression prevented premature keratinocyte differentiation.

## Materials and Methods

### Tissue culture

Pooled primary human keratinocytes from different juvenile donors were obtained from PromoCell and grown in a 1:1 mixture of KSF-M (Gibco) and Medium 154 for keratinocytes (Gibco). The media was supplemented with Human Keratinocyte Growth Supplement (HKGS), bovine pituitary extract (BPE), human epidermal growth factor (hEGF) and 0.5x Antibiotic-Antimycotic (all supplements obtained from Gibco). Cells were cultured at 37 °C in a humified chamber with 5% CO_2_. *In vitro* keratinocyte differentiation was induced by the addition of 1.2 mM calcium to the media and seeding cells at full confluency.

### RNA knockdown

siRNA pools of 11-30 different siRNAs were synthesized and obtained from siTools (Hannus *et al*, 2014). For siRNA-mediated knockdown, 5-6 million primary human keratinocytes were electroporated with 1 nmol siPools using the Amaxa human keratinocyte Nucleofector kit (Lonza) according to the manufacturer’s instructions and the T-018 program of the Amaxa Nucleofector II device (Lonza). After nucleofection, cells recovered for 24 h.

### Organotypic human epidermal tissue

For the generation of organotypic human epidermal tissue, 550,000 human keratinocytes nucleofected with siRNAs were seeded onto a devitalized dermal matrix and raised to the air-liquid interface to initiate stratification and differentiation, as described previously (Truong *et al*, 2006; Sen *et al*, 2010).

### Immunofluorescence and tissue analysis

Seven micrometer thick paraffin-embedded cross sections of human organotypic skin cultures were deparaffinized and rehydrated by two changes of xylene and washed with descending ethanol concentrations. For antigen retrieval, the cross sections were boiled twice in Demasking buffer (10 mM sodium citrate, pH 6.0, with 0.05% Tween 20) followed by blocking in PBS with 10% bovine calf serum (BCS) for 20 min at RT. Antibodies were diluted in PBS with 1% BCS and incubated with the sections for 1h. Primary antibodies included the following: Filaggrin (Santa Cruz, sc-66192) at 1:50 dilution and keratin 10 (NeoMarkers, MS-611-P) at 1:400 dilution. Alexa-555 conjugated goat anti-mouse (life technologies, A21422, 1:300) was used as secondary antibody diluted in PBS with 1% BCS. Unbound antibodies were washed off after one hour of incubation with PBS (three times, 5 min, RT). Slides were mounted in DAPI Fluoromount-G (Biozol). Finally, cross sections were analyzed with an Axiovert 200M fluorescence microscope and the AxioVision software (Carl Zeiss) or a BZ-X800 fluorescence microscope and the BZ-X800 Analyzer software (Keyence).

### qRT-PCR analysis

Total RNA from organotypic skin cultures was isolated with the RNeasy Plus mini kit (Qiagen) according to the manufacturer’s instructions. Total RNA from keratinocytes was isolated with TRIzol (Invitrogen) as stated by the manufacturer. Quantified by NanoDrop, 1 µg total RNA was subjected to reverse transcription with the iScript cDNA synthesis kit (Bio-Rad). In case of RNA isolated with TRIzol, genomic DNA was removed using DNAse I (Thermo Fisher Scientific) prior to cDNA synthesis. For qRT-PCR measurements, the Takyon Mix (Eurogentec) was used with the CFX96 Touch Real-time PCR Detection System (Bio-Rad). Samples were run at least in duplicates and normalized to ribosomal protein L32 mRNA according to the ΔΔC_t_-method (Livak & Schmittgen, 2001). The following primer sequences were used: EGR3_fwd: 5’- ACGCCAAGATCCACCTCAAG-3’, EGR3_rev: 5’-GGAAAAGTGGGGATCTGGGG-3’, Filaggrin_fwd: 5’-AAAGAGCTGAAGGAACTTCTGG-3’, Filaggrin_rev: 5’- AACCATATCTGGGTCATCTGG-3’, Keratin1_fwd: 5’- TGAGCTGAATCGTGTGATCC-3’, Keratin1_rev: 5’-CCAGGTCATTCAGCTTGTTC- 3’, L32_fwd: 5’-AGGCATTGACAACAGGGTTC-3’, L32_rev: 5’- GTTGCACATCAGCAGCACTT-3’, LCE1A_fwd: 5‘- GAAGCGGACTCTGCACCTAGAA-3‘, LCE1A_rev: 5‘- AGGAGACAGGTGAGGAGGAAATG-3‘, LCE6A_fwd: 5‘- GTCCTGATCTCTCCTCTCGTCT-3‘, LCE6A_rev: 5’- CAAGATTGCTGCTTCTGCTGT-3’, LINC00941_fwd: 5’- GACCTTTTCAGGCCAGCATT-3’, LINC00941_rev: 5’-ACAATCTGGATAGAGGGCTCA-3’, MTA2_fwd: 5’-TGTACCGGGTGGGAGATTAC- 3’, MTA2_rev: 5’-GCCTCCACATTTCCATTTGC-3’, SPRR4_fwd: 5’- AGCCTCCAAGAGCAAACAGA-3’, SPRR4_rev: 5’-GCAGGAGGAGATGTGAGAGG- 3’. Statistical significance was tested by an unpaired *t*-test and corrected for multiple testing after Bonferroni (*adj. *P*-value < 0.05, **adj. *P*-value < 0.01, ***adj. *P*-value < 0.001, and ns = not significant).

### *In vitro* RNA pull-down with subsequent protein detection

For *in vitro* RNA transcription of LINC00941 the Sp6 RNA Polymerase was used according to the manufacturer’s instructions with the exception that LINC00941 was labelled with biotin-16-UTP (Roche). 2 µg of biotin-16-labelled LINC00941 were incubated with protein lysate of 3*10^7^ keratinocytes. Protein lysate was prepared beforehand by collecting the cells in Lysis buffer (50 mM Tris, pH 7.5, 150 mM NaCl, 5% glycine, 1 mM DTT, 1 mM EDTA, 0.1 U/µl RiboLock RNase Inhibitor (Thermo Fisher Scientific), 1 mM AEBSF, 1x cOmplete Protease Inhibitor Cocktail (Roche)) and clearing the lysates for 15 min, full speed at 4 °C. After 16 h at 4 °C, 5 µl of MyOne Streptavidin C1 Dynabeads (Invitrogen) were added, the mixture was incubated for 1 h at RT and subjected to three wash cycles of 10 min, each using 500 µl Wash buffer (150 mM NaCl, 50 mM Tris, pH 7.5, 10% glycine, 5 µM EDTA, 0.5% Igepal and 1x cOmplete Protease Inhibitor Cocktail (Roche)). After the final wash, magnetic beads were resuspended in 12 µl 1x Laemmli buffer with 10% 2-mercaptoethanol following Western Blot analysis.

### Western Blot analysis

Protein lysates from calcium-induced differentiated keratinocytes were obtained by scraping cells into a suitable amount of Lysis buffer. Lysates were cleared for 15 min, full speed at 4 °C and the protein concentration was determined via Bradford assay. 30 µg of total protein were separated on a 10% SDS-PAGE gel and transferred onto the Amersham Hybond ECL membrane (Cytavia Life Science) by semi-dry blot. Blocking and antibody dilution was done in 5% milk powder in TBS-T. Primary antibodies included MTA2 (Abcam, ab8106) at 1:1,000 dilution and β-actin (Abcam, ab6276) at 1:10,000 dilution. Secondary antibodies utilized were IRDye 800 goat anti-rabbit (Licor Biosciences, 926-32211) and IRDye 680 goat anti-mouse (Licor Biosciences, 926-32220), both at a 1:15,000 dilution. Western Blots were analyzed with the Odyssey Infrared Imager (Licor Biosciences).

### Lentiviral-mediated LINC00941 overexpression

For HEK293T transfection, Lipofectamine 3000 (Thermo Fisher Scientific) was used according to the manufacturer’s instruction. The lentiviral transfer vector pLARTA-LINC00941 and packaging construct vectors (pUG-MDG and pCMV-ΔR8.91) were applied in equimolar quantities. Viral particles were harvested 48 h and 72 h after transfection. For lentiviral transduction, 50,000 keratinocytes per well were seeded in a 6-well plate the day before infection. The next day, an appropriate amount of viral particles diluted in keratinocyte medium and 5 µg/ml polybrene were added to the cells. Infection was accomplished in a centrifugation step at RT with 250 rcf for 1h. Afterwards, cells were recovered in keratinocyte medium for at least 24 h.

### Mass spectrometry (MS) analysis

MS analysis was essentially performed as previously described (Hoffmeister *et al*, 2023). Proteins were separated on a NuPAGE 4-12% Bis-Tris gel (Invitrogen) according to the manufacturer’s instructions and gel lanes were was cut into consecutive slices. The gel slices were washed with 50 mM NH_4_HCO_3_, 50mM NH_4_HCO_3_/acetonitrile (3/1) and 50mM NH_4_HCO_3_/acetonitrile (1/1) while shaking gently in an orbital shaker (VXR basic Vibrax, IKA). Gel pieces were lyophilized after shrinking by 100% acetonitrile. To block cysteines, reduction with DTT was carried out for 30 min at 57 °C followed by an alkylation step with iodoacetamide for 30 min at room temperature in the dark. Subsequently, gel slices were washed and lyophilized again as described above. Proteins were subjected to *in gel* tryptic digest overnight at 37 °C with approximately 2 µg trypsin per 100 µl gel volume (Trypsin Gold, mass spectrometry grade, Promega). Peptides were eluted twice with 100 mM NH_4_HCO_3_ followed by an additional extraction with 50 mM NH_4_HCO_3_ in 50% acetonitrile. Prior to LC-MS/MS analysis, combined eluates were lyophilized and reconstituted in 20 µl of 1% formic acid. Separation of peptides by reversed-phase chromatography was carried out on an UltiMate 3000 RSLCnano System (Thermo Scientific) which was equipped with a C18 Acclaim Pepmap100 preconcentration column (100µm i.D.x20mm, Thermo Fisher) in front of an Acclaim Pepmap100 C18 nano column (75 µm i.d. × 150 mm, Thermo Fisher). A linear gradient of 4% to 40% acetonitrile in 0.1% formic acid over 90 min was used to separate peptides at a flow rate of 300 nl/min. The LC-system was coupled on-line to a maXis plus UHR-QTOF System (Bruker Daltonics) via a CaptiveSpray nanoflow electrospray source (Bruker Daltonics). Data-dependent acquisition of MS/MS spectra by CID fragmentation was performed at a resolution of minimum 60000 for MS and MS/MS scans, respectively. The MS spectra rate of the precursor scan was 2 Hz processing a mass range between m/z 175 and m/z 2000. Via the Compass 1.7 acquisition and processing software (Bruker Daltonics) a dynamic method with a fixed cycle time of 3 s and a m/z dependent collision energy adjustment between 34 and 55 eV was applied. Raw data processing was performed in Data Analysis 4.2 (Bruker Daltonics), and Protein Scape 3.1.3 (Bruker Daltonics) in connection with Mascot 2.5.1 (Matrix Science) facilitated database searching of the Swiss-Prot *Homo sapiens* database (release-2020_01, 220420 entries). Search parameters were as follows: enzyme specificity trypsin with 1 missed cleavage allowed, precursor tolerance 0.02 Da, MS/MS tolerance 0.04 Da. Carbamidomethylation or propionamide modification of cysteine, oxidation of methionine, deamidation of asparagine and glutamine were set as variable modifications. Mascot peptide ion-score cut-off was set 25. Search conditions were adjusted to provide a false discovery (FDR) rate of less than 1%. Protein list compilation was done using the Protein Extractor function of Protein Scape. EmPAI-values (exponentially modified protein abundance index), which can be used for an approximate relative quantitation of proteins in a mixture, were extracted from Mascot. MS analysis resulted in *n* = 627 LINC00941 interaction partners. GO term analysis was carried out using the PANTHER Overrepresentation Test (Carbon & Mungall, 2023). GO terms with an FDR of less than 0.05 and chromatin association were considered for further analysis.

### RNA-Immunoprecipitation (RNA-IP)

For RNA-IP, cell lysate of *in vitro* differentiated keratinocytes was obtained by scraping cells into a suitable amount of RIPA buffer. Lysates were cleared for 10 min, full speed at 4 °C. 5 µg of either α-MTA2 antibody (Abcam, ab8106) or IgG control (Santa Cruz, I5006) were added to the lysate and incubate at 4 °C for 2 h. 40 µl Protein G Dynabeads (Invitrogen) were transferred to the lysate and incubated for 1 h at 4 °C. After three washing steps with RIPA buffer, the beads were resuspended in TRIzol (Invitrogen), and RNA was isolated.

### Chromatin Immunoprecipitation (ChIP) sequencing

For ChIP, primary keratinocytes were grown to a confluency of about 80% and dethatched from cell culture dishes. Following a cross-linking step adding 1% formaldehyde at RT for 10 min, the reaction was quenched with 125 mM glycine at RT for 5 min. Cells were pelleted and then lysed in Swelling buffer (100 mM Tris, pH 7.5, 10 mM potassium acetate, 15 mM magnesium acetate, 1% Igepal, 1x cOmplete Protease Inhibitor Cocktail (Roche) and 1 mM AEBSF). Chromatin fragmentation was subsequently performed in RIPA buffer for 30 min using the S220 Focused-ultrasonicator (Covaris) in the Freq sweeping mode, intensity 8, 20% duty cycle and 200 cycles per burst. 100 µg fragmented chromatin underwent IP with 5 µg α-MTA2 antibody (Abcam, ab8106) for 16 h at 4 °C. 40 µl Protein G Dynabeads (Invitrogen) were added to the IP and incubated for 45 min at RT. The beads were washed three times with Wash buffer (100 mM Tris, pH 9.0, 0.5 M lithium chloride, 1% Igepal, 1% sodium deoxycholate and 1 mM AEBSF) followed by the elution with Elution buffer (50 mM sodium bicarbonate and 1% SDS). For reverse cross-linking, 0.2 M sodium chloride was added, and the chromatin incubated at 67 °C for 4 h following a purification step with the QIAquick PCR Purification kit (Qiagen) according to the manufacturer’s instructions. For library preparation, the NEBNext Ultra II DNA Library Preparation kit for Illumina (New England Biolabs) and NEBNext Multiplex Oligos for Illumina (New England Biolabs) was used following the manufacturer’s instructions. The libraries were pooled equimolar, and the pool was quantified using the KAPA Library Quantification kit – Illumina (Roche). The libraries were sequenced on an Illumina NextSeq 2000 instrument (Illumina) controlled by the NextSeq Control Software (NCS) v1.4.1.39716, using a 100 cycles P2 Flow Cell with the single index, paired-end (PE) run parameters. Image analysis and base calling were done by the Real Time Analysis Software (RTA) v3.9.25. The resulting .cbcl files were converted into .fastq files with the bcl2fastq v2.20 software. ChIP sequencing was performed at the Genomics Core Facility “KFB – Center of Excellence for Fluorescent Bioanalytics” (University of Regensburg, Germany).

### ChIP sequencing data analysis

Initially, quality control of the raw sequence reads was conducted using FastQC (v0.11.8) (Andrews, 2010). Subsequently, reads were mapped to the reference genome (GRCh38) using bowtie2 (version v2.4.4). The following options were used to optimize the alignment process: --very-sensitive-local, --no-discordant, --no-mix, -- dovetail. Aligned reads were filtered for MAPQ >= 30 using samtools (Li *et al*, 2009). Reads mapping to blacklisted genomic regions (Encode accession: ENCFF356LFX) were removed using bedtools (Quinlan & Hall, 2010). Peak calling was executed on all samples relative to input samples using the MACS2 software (Zhang *et al*, 2008). The specific options used for this analysis were as follows: -f BAMPE, -g 2.7e9, --keep-dup auto. Following peak calling, the resultant peaks were filtered based on a log q-value threshold > 20 for subsequent analyses. This stringent cutoff was used to ensure a high level of confidence in the identified peaks, minimizing the potential for false positives. The number of fragments in each sample falling under these peaks was quantified using the featureCounts function of the Subread package (Liao *et al*, 2014).

To identify changes in MTA2 binding upon LINC00941 knock-down the generated count table was processed in R using the Bioconductor package DESeq2 (Love *et al*, 2014). 33 MTA2 binding sites were found to significantly change their binding abundance upon LINC00941 knock-down based on an FDR of 0.05.

### RNA sequencing data analysis

Publicly available RNA sequencing data of LINC00941 knock-down during keratinocyte differentiation (GSE118077) were re-analyzed using a nextflow RNA sequencing pipeline (Di Tommaso *et al*, 2017).

Initially, quality control of the raw sequence reads was conducted using FastQC (v0.11.8) (Andrews, 2010). Subsequently, reads were mapped to the reference genome (GRCh38) and corresponding gene annotation (Ensembl version 106) using the Spliced Transcripts Alignment to a Reference (STAR) software (version v2.7.8a) (Dobin *et al*, 2013). The following options were used to optimize the alignment process: --outFilterType BySJout, --outFilterMultimapNmax 20, --alignSJoverhangMin 8, -- alignSJDBoverhangMin 1, --outFilterMismatchNmax 999, --alignIntronMin 10, -- alignIntronMax 1000000, --outFilterMismatchNoverReadLmax 0.04, --runThreadN 12, --outSAMtype BAM SortedByCoordinate, --outSAMmultNmax 1, and -- outMultimapperOrder Random.

Post-mapping quality control was performed using the rnaseq mode of Qualimap (v2.2.1) (García-Alcalde *et al*, 2012). The level of PCR duplication was assessed using Picard MarkDuplicates (v2.21.8) (Broad Institute, 2019), and dupRadar (version v1.15.0) (Sayols *et al*, 2016). Gene expression quantification was carried out using featureCounts (v1.6.3) (Liao *et al*, 2014).

Finally, differential gene expression analysis between LINC00941 knockdown and control samples was conducted using the Bioconductor DESeq2 package (Love *et al*, 2014). The normal shrinkage method was applied for scaling log2 fold changes (Zhu *et al*, 2019). Genes with an FDR of less than 0.05 were considered significantly differentially expressed. This analysis resulted in the identification of 385 downregulated and 1532 upregulated genes in the LINC00941 knockdown keratinocytes at day 3.

### Downstream analysis

ChIP sequence coverage tracks were generated using the bamcoverage function from deeptools package (Ramírez *et al*, 2016). For normalization, the size factors derived from DESeq2 analysis were used. Data track visualizations were generated using the Bioconductor package Gviz (Hahne & Ivanek, 2016). The Bioconductor package ChIPseeker was used to annotate the genomic regions of the MTA2 binding sites (Yu *et al*, 2015). To link the MTA2 binding sites to the next gene only protein-coding genes were considered. Gene set over-representation analysis was carried out on MTA-associated genes with a maximum distance to the TSS of 3 kb using the Bioconductor package clusterProfiler (Wu *et al*, 2021).

### External data sets used in this study

The 15-state core chromatin model of male keratinocytes (roadmap accession: E057) was obtained from the roadmap repository (Kundaje *et al*, 2015) (https://egg2.wustl.edu/roadmap/web_portal/chr_state_learning.html). Corresponding Histone modification and RNA sequencing data tracks were obtained from ENCODE repository (Encode Project consortium, 2012; Luo *et al*, 2020): poly(A) RNA sequencing plus/minus strand (ENCFF283IQC/ENCFF804BRH), H3K4me3 (ENCFF517FHJ), H3K27me3 (ENCFF400FLX), H3K4me1 (ENCFF319BIJ). Data on *in vitro* differentiation of foreskin keratinocytes (at 0d/2.5d/5.5d) was obtained from the ENCODE Portal: ChromHMM 18-state model (ENCFF571LVQ/ENCFF385FJV/ENCFF058GJN), H3K4me1 (ENCFF539WZS/ENCFF546FWF/ENCFF821VJZ), H3K27ac (ENCFF786TBH/ENCFF870GTM/ENCFF988UOV), H3K27me3(ENCFF492WYD/ENCFF846IJG/ENCFF902OOG), poly(A) RNA sequencing plus (ENCFF646IJP/ENCFF786AVM/ENCFF533VFT), poly(A) RNA sequencing minus (ENCFF699EVT/ ENCFF701XBY/ENCFF366JLV), poly(A) RNA sequencing quantification (ENCFF423MWU/ ENCFF137YHI/ENCFF379PNP) RNA sequencing data of LINC00941 knock-down after day 2 and 3 was obtained from Gene Expression Omnibus (GEO; GSE118077) (Ziegler *et al*, 2019).

## Data availability

The datasets and computer code produced in this study are available in the following databases:

- MS data have been deposited to the ProteomeXchange Consortium via the PRIDE repository and will be released upon publication of this manuscript
- The code used to analyze the sequencing data: GitHub (https://github.com/uschwartz/linc00941_MTA2_kerationcytes)
- Raw and processed ChIP sequencing data have been deposited at GEO and will be released upon publication of this manuscript

## Acknowledgement

We thank Gernot Längst for critically reading this manuscript. Our research is supported by the Deutsche Forschungsgemeinschaft (SFB 960 to M.K.). We further thank the ENCODE Consortium for providing and the Manolis Kellis and Michael Snyder Lab for producing the datasets of human keratinocytes used in this study.

## Author contributions

EM JG, AB performed the experiments. EM, JG and MK designed the study. EM, US and MK analyzed the results. EM and MK drafted the manuscript.

---

**Supplement Figure 1:**
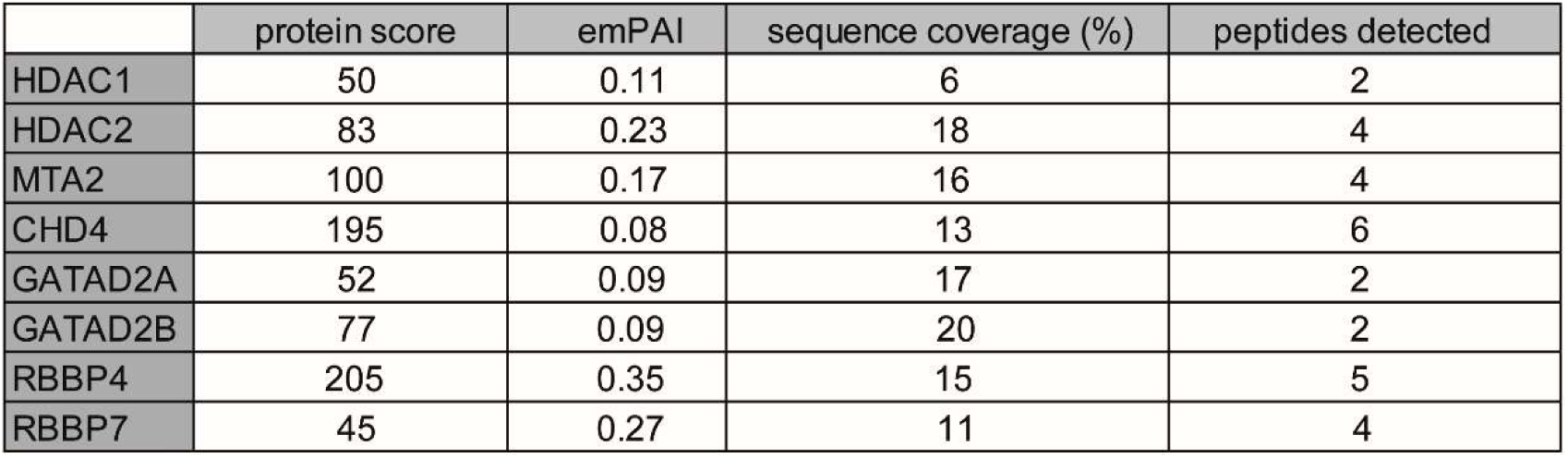
LINC00941 interacts with components of the NuRD complex. Quantitative analysis of NuRD components detected in MS analysis upon RNA-IP.

**Supplement Figure 3:**
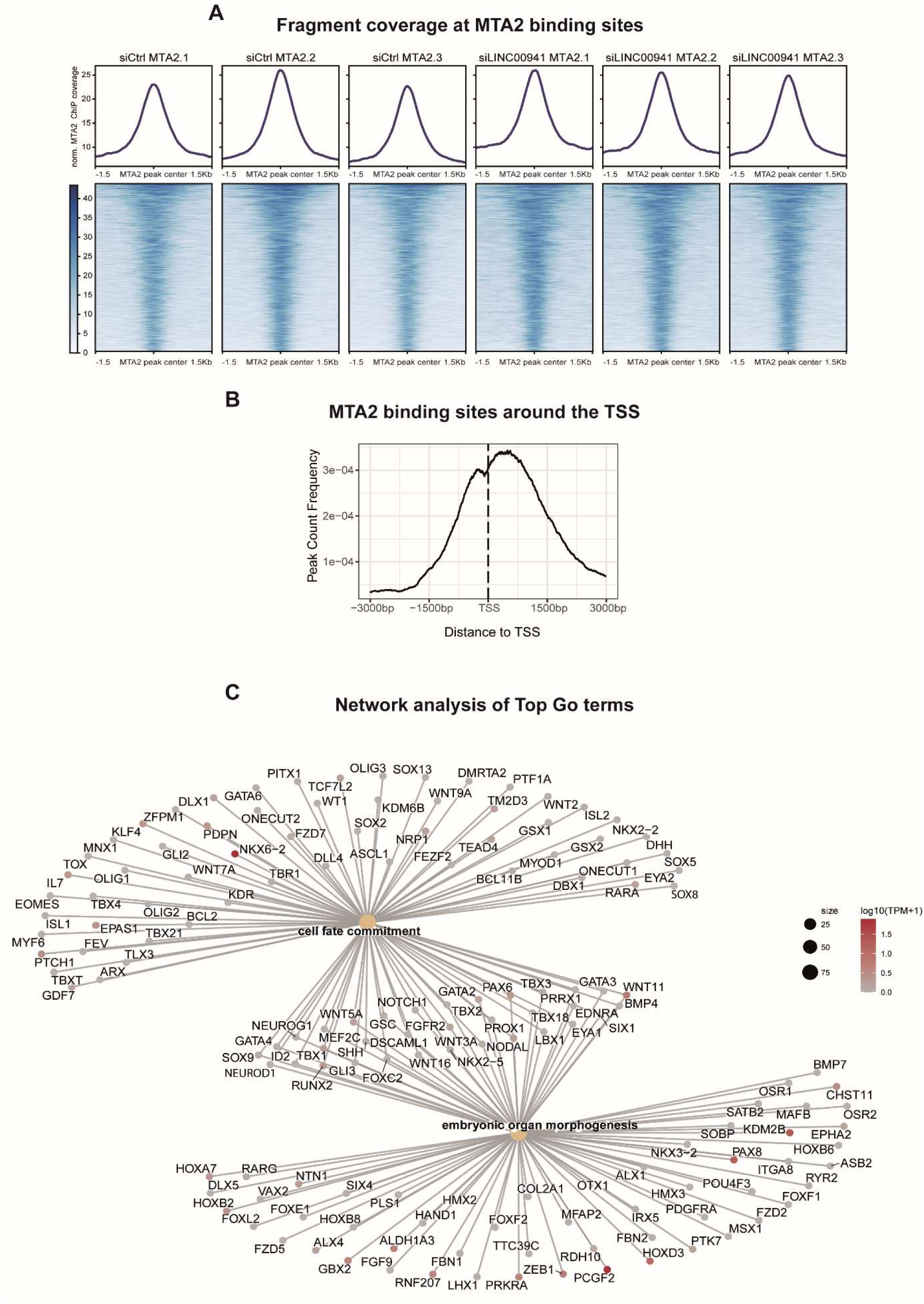
MTA2/NuRD occupies regulatory regions in keratinocytes. A) MTA2 ChIP sequencing fragment coverage at MTA2 binding sites of all replicates. The top panel shows the average fragment coverage of each sample across all called MTA2 peaks (*n* = 3,617). The bottom panel depicts the fragment coverage over the MTA2 bound sites as heatmap. B) Average profile of MTA2 binding sites around the TSS. C) Gene network plot of Top 2 GO terms of MTA2 ChIP sequencing including keratinocyte differentiation- and development-associated genes. The beige nodes symbolize enriched GO terms, with the size of the node indicating the number of associated genes. The lines connecting the nodes denote the specific genes associated with each GO term. The color of the gene nodes signifies the expression levels (log10(TPM+1)) in undifferentiated keratinocytes (ENCODE, ENCFF902PPD).

**Supplement Figure 4:**
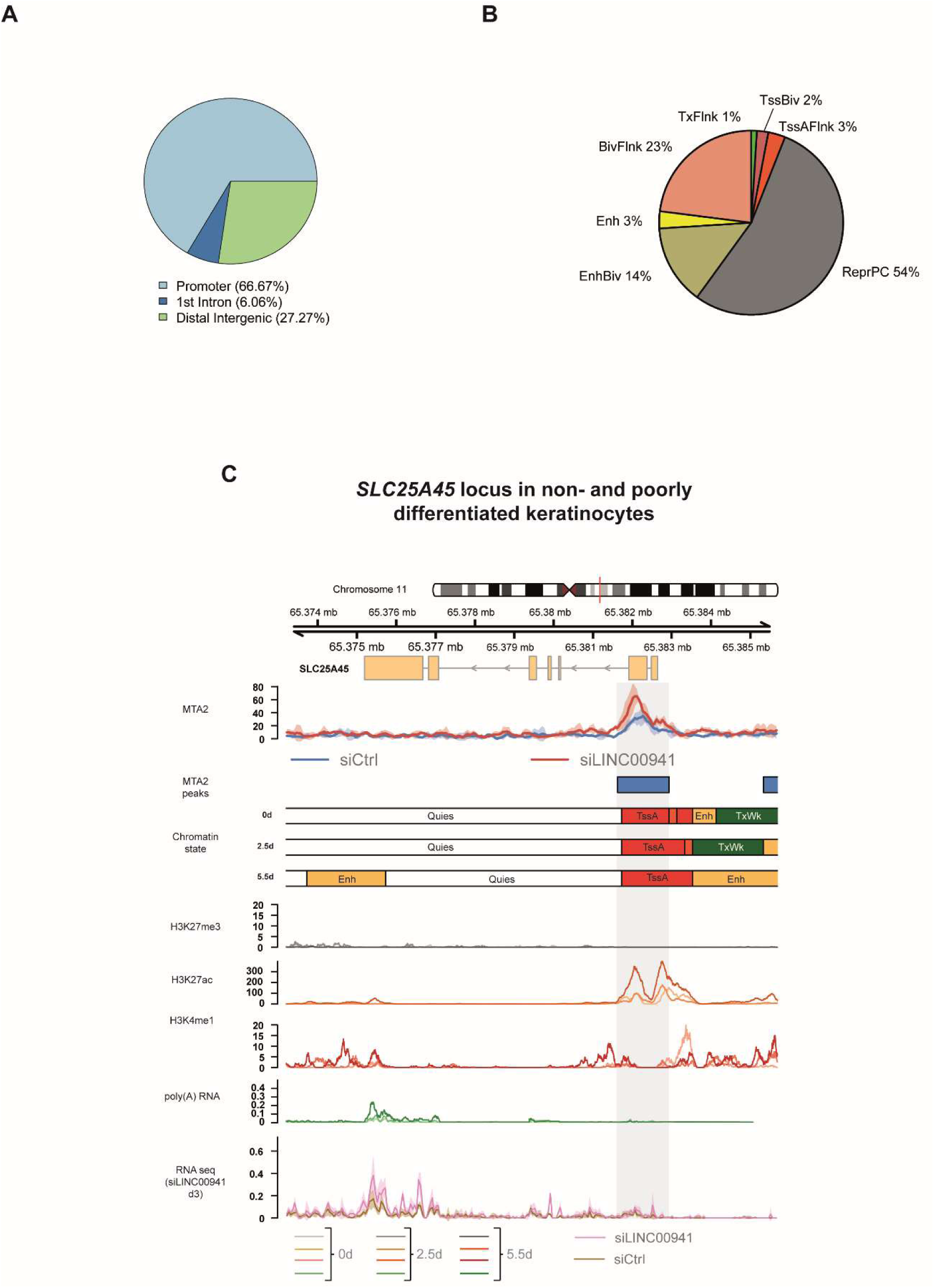
LINC00941 dependency of NuRD-associated MTA2 binding in *EGR3*. A, B) Pie chart showing the distribution of differential peaks at genomic features or chromatin states. LINC00941 knockdown altered MTA2/NuRD occupancy preferentially in promoter regions (A) mainly either annotated as repressed (ReprPC) or as bivalent (EnhBiv, BivFlnk) chromatin state (B) suggesting LINC00941 as putative regulator of transcription through MTA2/NuRD. The promoter region was defined as +/-1000 bp distance to the TSS of protein-coding genes. Abbreviations: Tss = Transcription Start Site; A = Active; Flnk = Flanking; Biv = Bivalent; Tx = Transcription; Enh = Enhancer; Repr = Repressed; PC = Polycomb; Quies = Quiescent C) Genome browser view of differential MTA2 binding sites at *SLC25A45* locus. Tracks of chromatin states, histone modifications, and transcription in primary undifferentiated (0d) and calcium-treated differentiated (2.5d, 5.5d) keratinocytes obtained from ENCODE portal are shown below the MTA2 ChIP sequencing tracks. Color shades in H3K27me3, H3K27ac, H3K4me1 and RNA sequencing tracks indicate the time points of differentiation: light shade for 0d, medium shade for 2.5d and dark shade for 5.5d. The bottom track shows the RNA sequencing coverage of siLINC00941 transfected (pink) and control (brown) keratinocytes after 3d of differentiation. The dashed box highlights the differential MTA2 binding site at *SLC25A45*. Abbreviations: Tss = Transcription Start Site; A = Active; Biv = Bivalent; Tx = Transcription; Wk = Weak; Enh = Enhancer; Quies = Quiescent

**Supplement Figure 5:**
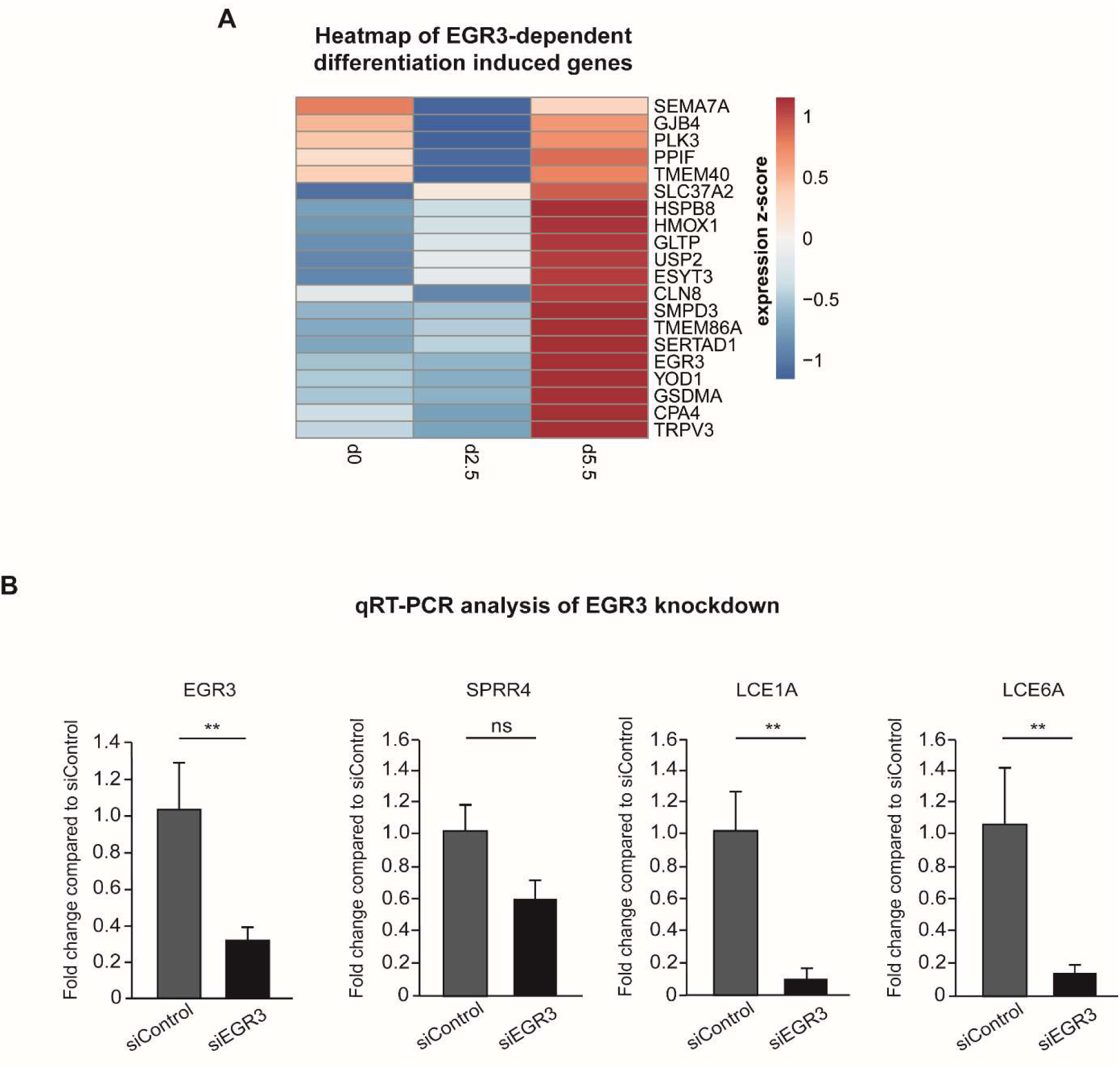
EGR3 induces differentiation of human organotypic epidermis. A) Heatmap showing transcriptional changes of EGR3 and EGR3-regulated genes in calcium-induced differentiated keratinocytes. TPM expression values after 0d, 2.5d and 5.5d were obtained from ENCODE Portal (ENCFF423MWU/ ENCFF137YHI/ENCFF379PNP) and transformed to z-scores. B) qRT-PCR analysis of EGR3 knockdown. SiPool-mediated knockdown of EGR3 in primary keratinocytes results in decreased abundance of EDC genes *SPRR4*, *LCE1A* and *LCE6A* on day 3 of differentiation in organotypic epidermal tissue (*n* = 3-4 tissue cultures/knockdown group). Data are presented as mean ± standard deviation. Statistical significance was tested by an unpaired *t*-test and corrected for multiple testing after Bonferroni (**adj. *P*-value < 0.01, and ns = not significant).

## References

Ai Y, Wu S, Zou C, Wei H (2020) LINC00941 promotes oral squamous cell carcinoma progression via activating CAPRIN2 and canonical WNT/β-catenin signaling pathway. J Cell Mol Med 24: 10512–10524

Andrews S (2010) FastQC: a quality control tool for high throughput sequence data. https://www.bioinformatics.babraham.ac.uk/projects/fastqc/

Arends T, Dege C, Bortnick A, Danhorn T, Knapp JR, Jia H, Harmacek L, Fleenor CJ, Straign D, Walton K et al (2019) CHD4 is essential for transcriptional repression and lineage progression in B lymphopoiesis. Proc Natl Acad Sci U S A 116: 10927–10936

Barrero MJ, Sese B, Kuebler B, Bilic J, Boue S, Martí M, Izpisua Belmonte JC (2013) Macrohistone variants preserve cell identity by preventing the gain of H3K4me2 during reprogramming to pluripotency. Cell Rep 3: 1005–1011

Bernstein BE, Mikkelsen TS, Xie X, Kamal M, Huebert DJ, Cuff J, Fry B, Meissner A, Wernig M, Plath K et al (2006) A bivalent chromatin structure marks key developmental genes in embryonic stem cells. Cell 125: 315–326

Blanpain C, Fuchs E (2006) Epidermal stem cells of the skin. Annu Rev Cell Dev Biol 22: 339–373

Blanpain C, Fuchs E (2009) Epidermal homeostasis: a balancing act of stem cells in the skin. Nat Rev Mol Cell Biol 10: 207–217

Bonasio R, Shiekhattar R (2014) Regulation of transcription by long noncoding RNAs. Annu Rev Genet 48: 433–455

Broad Institute (2019) Picard toolkit. https://Broadinstitute.Github.Io/Picard/

Cai P, Otten ABC, Cheng B, Ishii MA, Zhang W, Huang B, Qu K, Sun BK (2020) A genome-wide long noncoding RNA CRISPRi screen identifies PRANCR as a novel regulator of epidermal homeostasis. Genome Res 30: 22–34

Carbon S, Mungall C (2023) Gene Ontology Data Archive: Zenodo

Di Tommaso P, Chatzou M, Floden EW, Barja PP, Palumbo E, Notredame C (2017) Nextflow enables reproducible computational workflows. Nat Biotechnol 35: 316–319

Djebali S, Davis CA, Merkel A, Dobin A, Lassmann T, Mortazavi A, Tanzer A, Lagarde J, Lin W, Schlesinger F et al (2012) Landscape of transcription in human cells. Nature 489: 101–108

Dobin A, Davis CA, Schlesinger F, Drenkow J, Zaleski C, Jha S, Batut P, Chaisson M, Gingeras TR (2013) STAR: ultrafast universal RNA-seq aligner. Bioinformatics 29: 15– 21

Dubey N, Hoffman JF, Schuebel K, Yuan Q, Martinez PE, Nieman LK, Rubinow DR, Schmidt PJ, Goldman D (2017) The ESC/E(Z) complex, an effector of response to ovarian steroids, manifests an intrinsic difference in cells from women with premenstrual dysphoric disorder. Mol Psychiatry 22: 1172–1184

Encode Project consortium (2012) An integrated encyclopedia of DNA elements in the human genome. Nature 489: 57–74

García-Alcalde F, Okonechnikov K, Carbonell J, Cruz LM, Götz S, Tarazona S, Dopazo J, Meyer TF, Conesa A (2012) Qualimap: evaluating next-generation sequencing alignment data. Bioinformatics 28: 2678–2679

Hahne F, Ivanek R (2016) Visualizing Genomic Data Using Gviz and Bioconductor. Methods Mol Biol 1418: 335–351

Hannus M, Beitzinger M, Engelmann JC, Weickert M-T, Spang R, Hannus S, Meister G (2014) siPools: highly complex but accurately defined siRNA pools eliminate off-target effects. Nucleic Acids Res 42: 8049–8061

Harikumar A, Meshorer E (2015) Chromatin remodeling and bivalent histone modifications in embryonic stem cells. EMBO Rep 16: 1609–1619

Hoffmeister H, Holzinger S, Dürr M-S, Bruckmann A, Schindler S, Gröbner-Ferreira R, Depping R, Längst G (2023) Characterization of the nuclear import of human CHD4-NuRD complex. J Cell Sci

Hu G, Wade PA (2012) NuRD and pluripotency: a complex balancing act. Cell Stem Cell 10: 497–503

Kaji K, Caballero IM, MacLeod R, Nichols J, Wilson VA, Hendrich B (2006) The NuRD component Mbd3 is required for pluripotency of embryonic stem cells. Nat Cell Biol 8: 285–292

Kashiwagi M, Morgan BA, Georgopoulos K (2007) The chromatin remodeler Mi-2beta is required for establishment of the basal epidermis and normal differentiation of its progeny. Development 134: 1571–1582

Kazimierczyk M, Wrzesinski J (2021) Long Non-Coding RNA Epigenetics. Int J Mol Sci 22

Kim K-H, Son ED, Kim H-J, Lee SH, Bae I-H, Lee TR (2019) EGR3 Is a Late Epidermal Differentiation Regulator that Establishes the Skin-Specific Gene Network. J Invest Dermatol 139: 615–625

Kim TW, Kang B-H, Jang H, Kwak S, Shin J, Kim H, Lee S-E, Lee S-M, Lee J-H, Kim J-H et al (2015) Ctbp2 Modulates NuRD-Mediated Deacetylation of H3K27 and Facilitates PRC2-Mediated H3K27me3 in Active Embryonic Stem Cell Genes During Exit from Pluripotency. Stem Cells 33: 2442–2455

Kinkley S, Helmuth J, Polansky JK, Dunkel I, Gasparoni G, Fröhler S, Chen W, Walter J, Hamann A, Chung H-R (2016) reChIP-seq reveals widespread bivalency of H3K4me3 and H3K27me3 in CD4(+) memory T cells. Nat Commun 7: 12514

Kretz M, Siprashvili Z, Chu C, Webster DE, Zehnder A, Qu K, Lee CS, Flockhart RJ, Groff AF, Chow J et al (2013) Control of somatic tissue differentiation by the long non-coding RNA TINCR. Nature 493: 231–235

Kretz M, Webster DE, Flockhart RJ, Lee CS, Zehnder A, Lopez-Pajares V, Qu K, Zheng GXY, Chow J, Kim GE et al (2012) Suppression of progenitor differentiation requires the long noncoding RNA ANCR. Genes Dev 26: 338–343

Kundaje A, Meuleman W, Ernst J, Bilenky M, Yen A, Heravi-Moussavi A, Kheradpour P, Zhang Z, Wang J, Ziller MJ et al (2015) Integrative analysis of 111 reference human epigenomes. Nature 518: 317–330

Leboeuf M, Terrell A, Trivedi S, Sinha S, Epstein JA, Olson EN, Morrisey EE, Millar SE (2010) Hdac1 and Hdac2 act redundantly to control p63 and p53 functions in epidermal progenitor cells. Dev Cell 19: 807–818

Lee CS, Mah A, Aros CJ, Lopez-Pajares V, Bhaduri A, Webster DE, Kretz M, Khavari PA (2018) Cancer-Associated Long Noncoding RNA SMRT-2 Controls Epidermal Differentiation. J Invest Dermatol 138: 1445–1449

Li H, Handsaker B, Wysoker A, Fennell T, Ruan J, Homer N, Marth G, Abecasis G, Durbin R (2009) The Sequence Alignment/Map format and SAMtools. Bioinformatics 25: 2078–2079

Liao Y, Smyth GK, Shi W (2014) featureCounts: an efficient general purpose program for assigning sequence reads to genomic features. Bioinformatics 30: 923–930

Livak KJ, Schmittgen TD (2001) Analysis of relative gene expression data using real-time quantitative PCR and the 2(-Delta Delta C(T)) Method. Methods 25: 402–408

Love MI, Huber W, Anders S (2014) Moderated estimation of fold change and dispersion for RNA-seq data with DESeq2. Genome Biol 15: 550

Low JKK, Silva APG, Sharifi Tabar M, Torrado M, Webb SR, Parker BL, Sana M, Smits C, Schmidberger JW, Brillault L et al (2020) The Nucleosome Remodeling and Deacetylase Complex Has an Asymmetric, Dynamic, and Modular Architecture. Cell Rep 33: 108450

Lu J-T, Yan Z-Y, Xu T-X, Zhao F, Liu L, Li F, Guo W (2023) Reciprocal regulation of LINC00941 and SOX2 promotes progression of esophageal squamous cell carcinoma. Cell Death Dis 14: 72

Lu X, Chu C-S, Fang T, Rayon-Estrada V, Fang F, Patke A, Qian Y, Clarke SH, Melnick AM, Zhang Y et al (2019) MTA2/NuRD Regulates B Cell Development and Cooperates with OCA-B in Controlling the Pre-B to Immature B Cell Transition. Cell Rep 28: 472–485.e5

Lu X, Kovalev GI, Chang H, Kallin E, Knudsen G, Xia L, Mishra N, Ruiz P, Li E, Su L et al (2008) Inactivation of NuRD component Mta2 causes abnormal T cell activation and lupus-like autoimmune disease in mice. J Biol Chem 283: 13825–13833

Luo Y, Hitz BC, Gabdank I, Hilton JA, Kagda MS, Lam B, Myers Z, Sud P, Jou J, Lin K et al (2020) New developments on the Encyclopedia of DNA Elements (ENCODE) data portal. Nucleic Acids Res 48: D882–D889

Marques JG, Gryder BE, Pavlovic B, Chung Y, Ngo QA, Frommelt F, Gstaiger M, Song Y, Benischke K, Laubscher D et al (2020) NuRD subunit CHD4 regulates super-enhancer accessibility in rhabdomyosarcoma and represents a general tumor dependency. Elife 9

Miccio A, Wang Y, Hong W, Gregory GD, Wang H, Yu X, Choi JK, Shelat S, Tong W, Poncz M et al (2010) NuRD mediates activating and repressive functions of GATA-1 and FOG-1 during blood development. EMBO J 29: 442–456

Mikkelsen TS, Ku M, Jaffe DB, Issac B, Lieberman E, Giannoukos G, Alvarez P, Brockman W, Kim T-K, Koche RP et al (2007) Genome-wide maps of chromatin state in pluripotent and lineage-committed cells. Nature 448: 553–560

Morgenstern E, Kretz M (2023) The human long non-coding RNA LINC00941 and its modes of action in health and disease. Biol Chem

Pantier R, Mullin NP, Chambers I (2017) A new twist to Sin3 complexes in pluripotent cells. EMBO J 36: 2184–2186

Pundhir S, Su J, Tapia M, Hansen AM, Haile JS, Hansen K, Porse BT (2023) The impact of SWI/SNF and NuRD inactivation on gene expression is tightly coupled with levels of RNA polymerase II occupancy at promoters. Genome Res 33: 332–345

Quinlan AR, Hall IM (2010) BEDTools: a flexible suite of utilities for comparing genomic features. Bioinformatics 26: 841–842

Quinn JJ, Chang HY (2016) Unique features of long non-coding RNA biogenesis and function. Nat Rev Genet 17: 47–62

Ramírez F, Ryan DP, Grüning B, Bhardwaj V, Kilpert F, Richter AS, Heyne S, Dündar F, Manke T (2016) deepTools2: a next generation web server for deep-sequencing data analysis. Nucleic Acids Res 44: W160–5

Reynolds N, Salmon-Divon M, Dvinge H, Hynes-Allen A, Balasooriya G, Leaford D, Behrens A, Bertone P, Hendrich B (2012) NuRD-mediated deacetylation of H3K27 facilitates recruitment of Polycomb Repressive Complex 2 to direct gene repression. EMBO J 31: 593–605

Sanli I, Lalevée S, Cammisa M, Perrin A, Rage F, Llères D, Riccio A, Bertrand E, Feil R (2018) Meg3 Non-coding RNA Expression Controls Imprinting by Preventing Transcriptional Upregulation in cis. Cell Rep 23: 337–348

Sayols S, Scherzinger D, Klein H (2016) dupRadar: a Bioconductor package for the assessment of PCR artifacts in RNA-Seq data. BMC Bioinformatics 17: 428

Sen GL, Reuter JA, Webster DE, Zhu L, Khavari PA (2010) DNMT1 maintains progenitor function in self-renewing somatic tissue. Nature 463: 563–567

Strehle M, Guttman M (2020) Xist drives spatial compartmentalization of DNA and protein to orchestrate initiation and maintenance of X inactivation. Current Opinion in Cell Biology 64: 139–147

Tanis SEJ, Köksal ES, van Buggenum JAGL, Mulder KW (2019) BLNCR is a long non-coding RNA adjacent to integrin beta-1 that is rapidly lost during epidermal progenitor cell differentiation. Sci Rep 9: 31

Truong AB, Kretz M, Ridky TW, Kimmel R, Khavari PA (2006) p63 regulates proliferation and differentiation of developmentally mature keratinocytes. Genes Dev 20: 3185–3197

Ullah I, Thölken C, Zhong Y, John M, Rossbach O, Lenz J, Gößringer M, Nist A, Albert L, Stiewe T et al (2022) RNA inhibits dMi-2/CHD4 chromatin binding and nucleosome remodeling. Cell Rep 39: 110895

Voigt P, Tee W-W, Reinberg D (2013) A double take on bivalent promoters. Genes Dev 27: 1318–1338

Wu T, Hu E, Xu S, Chen M, Guo P, Dai Z, Feng T, Zhou L, Tang W, Zhan L et al (2021) clusterProfiler 4.0: A universal enrichment tool for interpreting omics data. Innovation (Camb) 2: 100141

Yildirim O, Li R, Hung J-H, Chen PB, Dong X, Ee L-S, Weng Z, Rando OJ, Fazzio TG (2011) Mbd3/NURD complex regulates expression of 5-hydroxymethylcytosine marked genes in embryonic stem cells. Cell 147: 1498–1510

Yu G, Wang L-G, He Q-Y (2015) ChIPseeker: an R/Bioconductor package for ChIP peak annotation, comparison and visualization. Bioinformatics 31: 2382–2383

Zahid H, Buchholz CR, Singh M, Ciccone MF, Chan A, Nithianantham S, Shi K, Aihara H, Fischer M, Schönbrunn E et al (2021) New Design Rules for Developing Potent Cell-Active Inhibitors of the Nucleosome Remodeling Factor (NURF) via BPTF Bromodomain Inhibition. J Med Chem 64: 13902–13917

Zhang Y, Liu T, Meyer CA, Eeckhoute J, Johnson DS, Bernstein BE, Nusbaum C, Myers RM, Brown M, Li W et al (2008) Model-based analysis of ChIP-Seq (MACS). Genome Biol 9: R137

Zhao Z, Sentürk N, Song C, Grummt I (2018) lncRNA PAPAS tethered to the rDNA enhancer recruits hypophosphorylated CHD4/NuRD to repress rRNA synthesis at elevated temperatures. Genes Dev 32: 836–848

Zhu A, Ibrahim JG, Love MI (2019) Heavy-tailed prior distributions for sequence count data: removing the noise and preserving large differences. Bioinformatics 35: 2084– 2092

Ziegler C, Graf J, Faderl S, Schedlbauer J, Strieder N, Förstl B, Spang R, Bruckmann A, Merkl R, Hombach S et al (2019) The long non-coding RNA LINC00941 and SPRR5 are novel regulators of human epidermal homeostasis. EMBO Rep 20

